# Distinct spectral profiles of awake resting EEG in disorders of consciousness - the role of frequency and topography of oscillations

**DOI:** 10.1101/2022.11.03.514999

**Authors:** Dominika Drążyk, Karol Przewrocki, Urszula Górska, Marek Binder

## Abstract

The prolonged disorders of consciousness (PDOC) pose a challenge for an accurate clinical diagnosis, mainly due to patients’ scarce or ambiguous behavioral responsiveness. Measurement of brain activity can support better diagnosis, independent of motor restrictions. Methods based on spectral analysis of resting-state EEG appear as a promising path, revealing specific changes within the internal brain dynamics in PDOC patients. In this study we used a robust method of resting-state EEG power spectrum parameter extraction to identify distinct spectral properties for different types of PDOC.

Sixty patients and 37 healthy volunteers participated in this study. Patient group consisted of 22 unresponsive wakefulness patients, 25 minimally conscious patients and 13 patients emerging from the minimally conscious state. Ten minutes of resting EEG was acquired during wakefulness and transformed into individual power spectra. For each patient, using the spectral decomposition algorithm, we extracted maximum peak frequency within 1-14 Hz range in the centro-parietal region, and the antero-posterior (AP) gradient of the maximal frequency peak. All patients were behaviorally diagnosed using Coma Recovery Scale - Revised (CRS-R).

The maximal peak frequency in the 1-14 Hz range successfully predicted both neurobehavioral capacity of patients as indicated by CRS-R total score and PDOC diagnosis. Additionally in patients in whom only one peak within the 1-14 Hz range was observed, the AP gradient significantly contributed to the accuracy of prediction. We have identified three distinct spectral profiles of patients, likely representing separate neurophysiological modes of thalamocortical functioning. Etiology did not have significant influence on the obtained results.

## Introduction

The prolonged disorders of consciousness (PDOC) are characterized by a pathological dissociation between *wakefulness* - an ability to orient toward and respond to stimuli, maintained by subcortical regions (Demertzi et al 2009), and *conscious awareness*, probably supported by the function of the thalamocortical system (Laureys 2005; Schiff 2010). Severe brain injuries, most frequently caused by trauma, stroke, or anoxia, may result in a state of coma (Posner et al 2019) that may further evolve into other forms of PDOC. These include *Unresponsive Wakefulness Syndrome* (UWS, Laureys et al 2010), which is characterized by the lack of self-awareness or the awareness of environment (Monti et al 2010), with preserved irregular sleep cycles as well as intact respiration and thermoregulatory capacity (Gosseries et al 2011). The recovery from UWS may result in a *Minimally Conscious State* (MCS), a state in which the patient is able to follow simple commands, visually pursuit objects and people or respond verbally (Schnakers et al 2016). However, those behavioral manifestations of consciousness may be intermittent in MCS. Reappearance of functional communication and/or ability to use objects adequately is a sign of the *Emergence from Minimally Conscious State* (EMCS, Giacino et al 2002). The correct differentiation between the above mentioned states remains a great challenge for medical personnel as well as caregivers. The repeatedly observed high rate of misdiagnosis of UWS (Childs et al 1993; Schnakers et al 2009; Forgacs et al 2014) only emphasizes the need for improved diagnostic protocols.

Standardized clinical tools, of which the gold standard is *Coma Recovery Scale-Revised* (CRS-R, Giacino et al 2004), may reduce the likelihood of diagnostic errors. Unfortunately however, their reliability is limited as they are based on the overt motor responses to sensory stimulation and verbal communication. Using them as the only diagnostic tool may result in the understatement of patients neurocognitive capacity, especially in cases of *Cognitive-Motor Dissociation* (CMD, Schiff et al 2014; Stender et al 2014; Schnakers 2020) or *Locked-In Syndrome* (LIS, Gosseries et al 2016), where motor unresponsiveness prevents patients from following commands.

The non-invasive, relatively low-cost and widely available EEG technique is a promising alternative that may contribute to a better diagnosis. However, active cognitive paradigms, e.g. motor imagery (Cruse et al 2011), may lead to false-negatives, due to the patient’s impaired speech comprehension, or limited motor capacity. In turn, passive EEG paradigms do not require volitional activity (Schnakers 2020). Recent approaches involve, for example, sensory evoked responses (Kotchoubey et al 2015; Binder et al 2017; Górska and Binder 2019), or TMS-EEG procedures (Casarotto et al 2016), but specifically the measurement of spontaneous, resting-state EEG signal can be regarded as a rich source of information about the functional integrity of the cortical mantle (Forgacs et al 2014). Frequency domain EEG analysis, representing contributions of individual frequencies in the signal, has already been proved very useful in PDOC diagnosis (Bai et al 2017).

It has been previously demonstrated that the ratio of high (8 - 30 Hz) to low (2 - 8 Hz) frequencies correlates with the CRS-R total score (Lechinger et al 2013). Other studies pointed out that the power of delta range (1-4 Hz) is related to bad outcome in PDOC patients (Lehembre et al 2012; Piarulli et al 2016). A good discriminative potential regarding the PDOC diagnosis was also assigned to the *slowing* of oscillatory power within the range of 4-13 Hz in the parietal region (Schiff et al 2014). Unfortunately, the mechanisms of this phenomenon are not well understood. In most cases EEG data is analyzed by averaging the amplitude within separate frequency bands. Therefore, it is not clear whether the observed effect is related to the shift of a single frequency peak related to increasing level of neurocognitive capacity or, alternatively, it is a result of independent low-frequency oscillatory activity in UWS and middle-frequency alpha-like activity in MCS and EMCS.

Previous studies also investigated differences in spatial dynamics of the patient’s spectral profile. For example, the antero-posterior (AP) gradient was used, representing the gradual increase in frequency with the parallel decrease in amplitude observed on the antero-posterior axis of a scalp, including dominant frontal beta activity (Forgacs et al 2017; Schiff et al 2014; Estraneo et al 2016). This feature illustrates characteristic dynamic differences when it comes to the distribution of alpha rhythm in wakefulness and sleep stages (DeGennaro et al 2001) and was shown as a promising feature of preserved corticothalamic integrity in PDOC patients (Forgacs et al 2017).

In this study we investigated the relationship between the frequency peaks of the highest amplitude within the 1-14 Hz range of the frequency spectra and the outcome of the CRS-R behavioral scale in the group of PDOC patients with various etiologies. Moreover, we examined the spatial dynamics of this oscillation, and evaluated the usefulness of AP gradient, previously advocated as a marker of the functional recovery in the PDOC group (Forgacs et al 2014, 2017; Schiff et al 2014; Estraneo et al 2016). While the existence and features of AP gradient were previously assessed by the expert qualitative evaluation, this research aims to investigate the possibility of its quantitative operationalization, with a use of FOOOF algorithm (*Fitting Oscillations & One Over F*, Donoghue et al 2020), a robust and easy-to-use spectral decomposition method. We found that the increase in the dominant peak predicts the increase of the CRS-R total score, while the outcome from the gradient measure depends on the integrity of the dominant peak in the EEG spectrum.

## Methods

### Subjects

The initial group of healthy controls (HC) included 45 subjects, each with the single EEG measurement. From the initial group 8 subjects were excluded due to the large muscle artifacts and/or an extensive noise in the EEG signal. Thus, the final dataset used in the analysis included 37 measurements obtained from HC (*N* = 37 including 22 females, aged 23.94 ± 3.15).

The initial group of PDOC patients comprised 134 measurements, obtained from 78 patients. Due to the large muscle artifacts and/or an extensive noise present in the EEG centro-parietal electrodes we excluded 49 measurements (see Tab. **1** for details). This amount, although high (~37%), presents no deviation from the common exclusion criteria in the area of research on patients with severe brain injuries (Cao et al 2019; Pistoia et al 2014; Kremneva et al 2019; Forgacs et al 2017). Finally, the analysis included a set of 86 measurements from 60 PDOC patients (*N* = 53, including 20 females, aged 36.88 ± 13.53, see Tab. **1** for details).

**Table 1.** Demographic features of groups of patients within a CRS-based diagnosis.

| Groups | N measurements | CRS-based diagnosis |  |  |  |
| --- | --- | --- | --- | --- | --- |
|  |  | N | Females | Age |  |
|  |  |  |  | M | SD |
| UWS | 30 | 22 | 5 | 36.33 | 12.43 |
| MCS | 36 | 25 | 12 | 39.31 | 15.47 |
| EMCS | 20 | 13 | 7 | 32.56 | 10.01 |

Table A1 in Appendix A (Supplementary information) provides detailed information about the PDOC patients, including the sex, age, etiology, CRS-R results with the subscales and diagnosis, time since injury in months, dates of EEG and CRS measurements, list of administered medications, and finally the exclusion criteria, when applicable.

An informed consent was obtained from each healthy participant and from the legal surrogates of all patients. The study was approved by the ethics review board of the Institute of Psychology, Jagiellonian University and was conducted in accordance with the Declaration of Helsinki (1975, revised 2000).

### EEG measurement

PDOC resting state EEG data was collected in three specialized in-patient centers for care and treatment in Torun, Czestochowa and Krakow, Poland. Healthy control group data were collected in the Psychophysiology Laboratory of the Institute of Psychology, Jagiellonian University in Krakow, Poland. The recording was performed using 64-electrode Active Two (BioSemi, Amsterdam, NL), with a 10–20 system headcap and four additional electrodes located above and below the right eye and in the external canthi of both eyes. Two additional reference electrodes were placed on mastoids and recorded in parallel. ‘CMS’ (*common mode sense*) and ‘DRL’ (*driven right leg*) electrodes were placed between ‘POz’ and ‘PO3’ and ‘POz’ and ‘PO4’, respectively.

Each time recording was carried on for 10 minutes and sampled at 1024 Hz. Patients were examined in the isolated room, reclining in bed or sitting on the wheelchair in a comfortable position, with their eyes open. Healthy control group was examined when sitting on a chair in the sound isolated laboratory room, with their gaze set on the fixation cross in the center of the computer screen, and were instructed not to focus on anything specific during the recording time.

### Preprocessing

The off-line preprocessing of EEG data was performed with Brain Vision Analyzer 2.0 software (Brain Products, Gilching). Noisy channels were interpolated with a spline method, data was filtered in 1-50 Hz range (IIR Filter, Zero phase shift Butterworth filter, order 8), and downsampled to 256 Hz. Raw data inspection excluded massive artifacts before the segmentation, whereas blink and eye movement artifacts were corrected using Infomax Independent Component Analysis (ICA). Data was further re-referenced to the common average. Next, signal from each channel was segmented into non-overlapping 2 s intervals, with the noisy segments excluded from the analysis using semi-automatic mode with the following criteria: amplitude limits −150 *μ*V to 150 *μ*V; 150 *μ*V maximum allowed difference in intervals over 200 ms; maximal voltage step of 75 *μ*V/ms. Fast Fourier Transform (FFT, 10% Hanning window, 0.5 Hz resolution) was calculated on the remaining segments and averaged, for each channel separately.

Due to frequent excessive movements, posturing and muscle twitches and structural brain damage, EEG data acquired from PDOC patients may become noisy, yet the signal from parietal and central regions of the scalp is usually the least affected by those artifacts (Forgacs et al 2020). Thus, the *centro-parietal region* channel, constructed by averaging the electrodes ‘Cz’, ‘CPz’, ‘Pz’, ‘CP3’, ‘CP1’, ‘P3’, ‘P1’, ‘CP2’, CP4’, ‘P2’, and ‘P4’, was used to calculate the frequency of the highest peak in the EEG spectrum and the ‘POz’, ‘Pz’ and ‘CPz’ electrodes were used to calculate *midline AP gradient*.

### Data analysis

The clinical state of the patients was determined using the Polish adaptation of the *Coma Recovery Scale - Revised* (CRS-R, Binder et al 2018). CRS-R is the most recommended clinical tool for behavioral assessment of neurocognitive functions in PDOC patients (Giacino et al 2009; Schnakers 2020). It consists of 23 items hierarchically arranged into six subscales that address auditory, visual, motor, oromotor, communication and arousal functions, so that within each subscale the lowest score corresponds to the reflexive responses, and the highest score indicates the presence of a cognitively mediated behavior (Giacino et al 2004). The experimenters were blind to the results of the CRS-R assessment at the time of recording and data preprocessing.

It should be also noted that in this experiment the CRS-R assessment was conducted only once, while recently its multiple administration was recommended in order to reduce the level of ambiguity in assessing the clinical condition of patients (Wannez et al 2017). Thus, it can not be ruled out that its single usage might have weakened the estimates of the existing relationship between the clinical state of the patient and the measured spectral parameters. Moreover, we were not able to administer the CRS-R test on exactly the same day as the EEG measurement every time (the difference between CRS-R and EEG measurements in days is 1.91 ± 3.96). Though the result of CRS-R assessment tends to change over longer time periods, it may be expected to remain stable over several days (Beukema et al 2016; Schiff et al 2007).

In the analysis described below, the total score of the CRS-R scale obtained for each patient is represented with the *CRSscore* variable, while the PDOC diagnosis obtained on the basis of individual CRS-R results (UWS, MCS or EMCS) with the *CRSdiagnosis* variable.

The averaged EEG spectra were parameterized using FOOOF algorithm (version 0.1.3) with the *fooof_mat* wrapper (version 0.0.1, Donoghue et al 2020) within the MATLAB environment (Mathworks Inc., version 2019b). The method allows for a precise isolation of frequency peaks in the EEG signal spectrum, while controlling for the background aperiodic signal component. It decomposes the EEG spectrum into an exponential two-parameter aperiodic component as well as a series of Gaussian peaks with parameters describing their center frequency, amplitude and bandwidth. The goodness of fit metrics are represented by *R*^2^ of the model fit and an error estimate. The settings for the algorithm were established as: peak width limits: 2-6; max number of peaks: 4; minimum peak height: 0.111; peak threshold: 2 and aperiodic mode: fixed. Power spectra were parameterized across frequency range between 1 to 45 Hz.

Based on FOOOF results, for each measurement we obtained a *MaxPeakFreq* parameter, which was the center frequency of the peak with the highest amplitude within the 1-14 Hz search range (extending across delta, theta and alpha EEG frequency bands) in the averaged signal from the centro-parietal region. Of note, differently from Forgacs et al. (2017) we focused the analysis only on the 1-14 Hz frequency range, as the noise level in the beta range (mainly consisting of muscle activity) did not allow us to draw reliable conclusions.

The *Gradient* parameter, representing the antero-posterior (AP) gradient of the amplitude of the highest peak with the central frequency in range 1-14 Hz, was calculated as follows: for the 1-14 Hz range of modeled power spectrum of each of the midline electrodes ‘POz’, ‘Pz’ and ‘CPz’, we found the amplitude of the highest peak. To make sure we are tracking the same peak, as it has been determined by the *MaxPeakFreq*, every other peak deviating from the *MaxPeakFreq* by at least half of its bandwidth on each side of the spectrum was rejected. For further analysis we used only those measurements in which the specified peak was successfully found in at least 2 of 3 midline electrodes, with at least 0.95 goodness of modeled power spectrum fit. Then we constructed a vector of peaks amplitudes, normalized by dividing each value by the maximal value of the vector. Finally, to obtain *Gradient* parameter we calculated vectors gradient, using *gradient* function (MATLAB version R2014b) and averaged it across three previously selected channels.

### Statistical analysis

To capture the relationship between the resting-state EEG measures and the patient neurobehavioral condition assessed using CRS-R scale, we introduced two separate models: *Model 1* with a total CRS-R score (labeled further as *CRSscore*) and *Model 2* with a CRS-R based diagnosis (labeled further as *CRSdiagnosis*), as respective outcomes. We adopted the total CRS-R score (range 2-23 with the mean of 10.714 ± 5.941), as the index of the overall level of neurocognitive capacity in PDOC patients (Bodart et al 2018; Rosanova et al 2012). The CRS-R-based PDOC diagnosis was treated as an ordinal categorical variable. This variable consisted of four ordered categories: from the group with the lowest (UWS), through intermediate (MCS) and the highest capacity (EMCS), to the healthy control group (HC). The HC group was included to illustrate the contrast between the patterns of resting state activity observed in an intact, aware brain and those with injuries affecting the state of consciousness.

#### Model 1

In the *CRSscore* analysis we decided to follow the multilevel models strategy, as it takes into account subject-level variability better than the traditional methods of data aggregation (Moscatelli et al 2012). Statistical analysis was performed using the R environment (R Core Team, 2021, version 4.0.3) with lme4 library (version 1.1-27.1, Bates et al 2006). LMMs were fitted using the maximum likelihood method (Laplace Approximation, Bates et al 2006). The structure of factors used in the model was constructed step up, from the model with Subject variable as a random intercept (*Model 1n*), to more complex ones with singular (*Model 1a* with *MaxPeakFreq*, *Model 1b* with *Gradient*) and both fixed factors (*Model 1c*) included. Random intercept structure used in every model controlled for the variable number of measurements obtained from each subject. Log-likelihood (*LL*) as an indicator of model selection was compared for all models, each time contrasting the model with the effect in question against the model without the effect in question. The one with the lowest log-likelihood value was considered final (Snijders and Bosker 2012).

#### Model 2

Probability of the given diagnosis (*CRSdiagnosis*) based on the *MaxPeakFreq* and *Gradient* was calculated using cumulative link mixed models (CLMM, Christensen 2018) for the ordinal outcomes with probit distribution. Analysis was performed using the ordinal library (version 2020.8-22, Bürkner 2017; Bürkner and Vuorre 2019). The structure of factors used in the model was constructed step up, from the model with Subject variable as a random intercept (*Model 2n*), to more complex models with single (*Model 2a* with *MaxPeakFreq*, *Model 2b* with *Gradient*) and both fixed factors (*Model 2c*) included. Random effects structure and model selection process was the same as in *CRSscore* analysis.

## Results

### Total CRS-R score in relation to the frequency of the maximum peak in the spectrum

Total CRS-R scores for individual patients are presented in Table **2** and the averaged spectra fitted using FOOOF parameters in Figure **1**. *Model 1a* was used to assess the relationship between *CRSscore* and the frequency of the highest peak in the 1-14 Hz range (*MaxPeakFreq*). Model presented better fit compared to the null model (Δ*LL* = −5.11, see Tab. **2**). Analysis showed a statistically significant effect of *MaxPeakFreq β* = 0.74 ± 0.22, *CI* : [0.30 1.20], see Fig. **2**A). On the contrary, *Model 1b* (*LL* = −251.39) with the antero-posterior gradient (*Gradient*) as a variable did not present a significant effect (*CI* : [-5.19 4.65], Fig. **2**B). Although *Model 1c* including the interaction of *MaxPeakFreq* and *Gradient* was characterized with minimally better fit (*LL* = −245.69) compared to the *Model 1a*, the improvement was too small to account for the cost of increased complexity of the model. The interaction itself was not significant (*CI* : [-1.63 2.13]), *Model 1c* was therefore rejected. Results indicate that the increase of *CRSscore* could be related to the increase of the frequency of the highest peak in the 1-14 Hz range, but not to the value of the antero-posterior gradient.

**Table 2.**
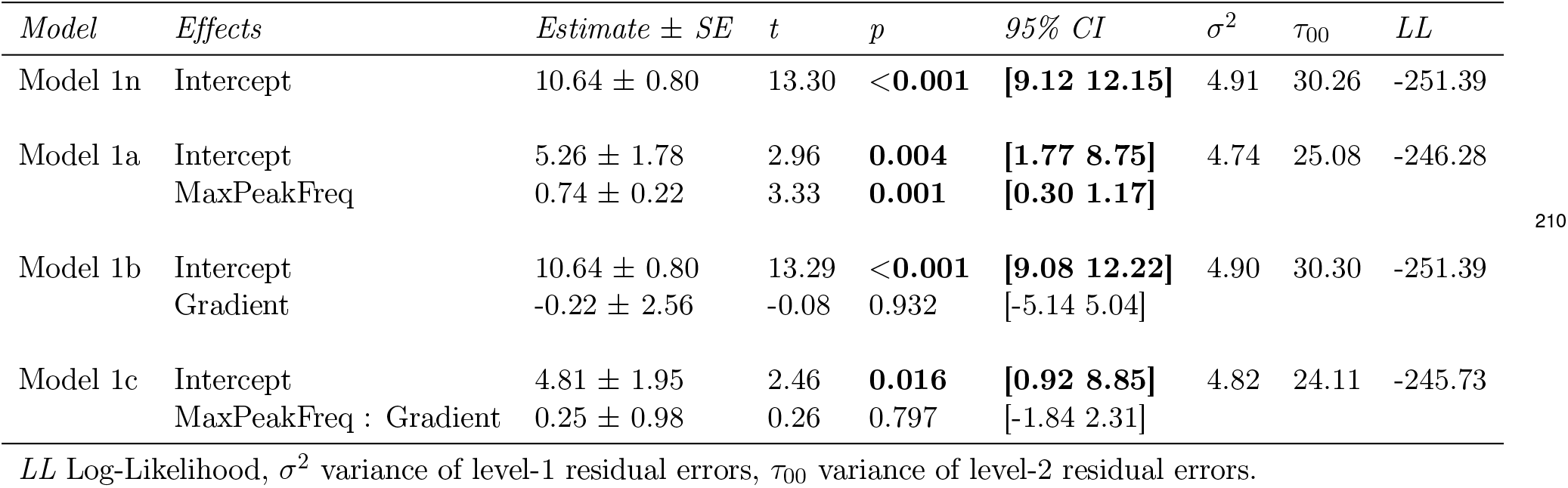
Summary of the *CRSscore* models fitted with its accuracy diagnostic.

| Model | Effects | Estimate ± SE | t | p | 95% CI | $\sigma^2$ | $\tau_{00}$ | LL |
| --- | --- | --- | --- | --- | --- | --- | --- | --- |
| Model 1n | Intercept | 10.64 ± 0.80 | 13.30 | <0.001 | [9.12 12.15] | 4.91 | 30.26 | -251.39 |
| Model 1a | Intercept | 5.26 ± 1.78 | 2.96 | 0.004 | [1.77 8.75] | 4.74 | 25.08 | -246.28 |
| | MaxPeakFreq | 0.74 ± 0.22 | 3.33 | 0.001 | [0.30 1.17] | | | |
| Model 1b | Intercept | 10.64 ± 0.80 | 13.29 | <0.001 | [9.08 12.22] | 4.90 | 30.30 | -251.39 |
| | Gradient | -0.22 ± 2.56 | -0.08 | 0.932 | [-5.14 5.04] | | | |
| Model 1c | Intercept | 4.81 ± 1.95 | 2.46 | 0.016 | [0.92 8.85] | 4.82 | 24.11 | -245.73 |
| | MaxPeakFreq : Gradient | 0.25 ± 0.98 | 0.26 | 0.797 | [-1.84 2.31] | | | |

*LL* Log-Likelihood, $\sigma^2$ variance of level-1 residual errors, $\tau_{00}$ variance of level-2 residual errors.

**Figure 1.**
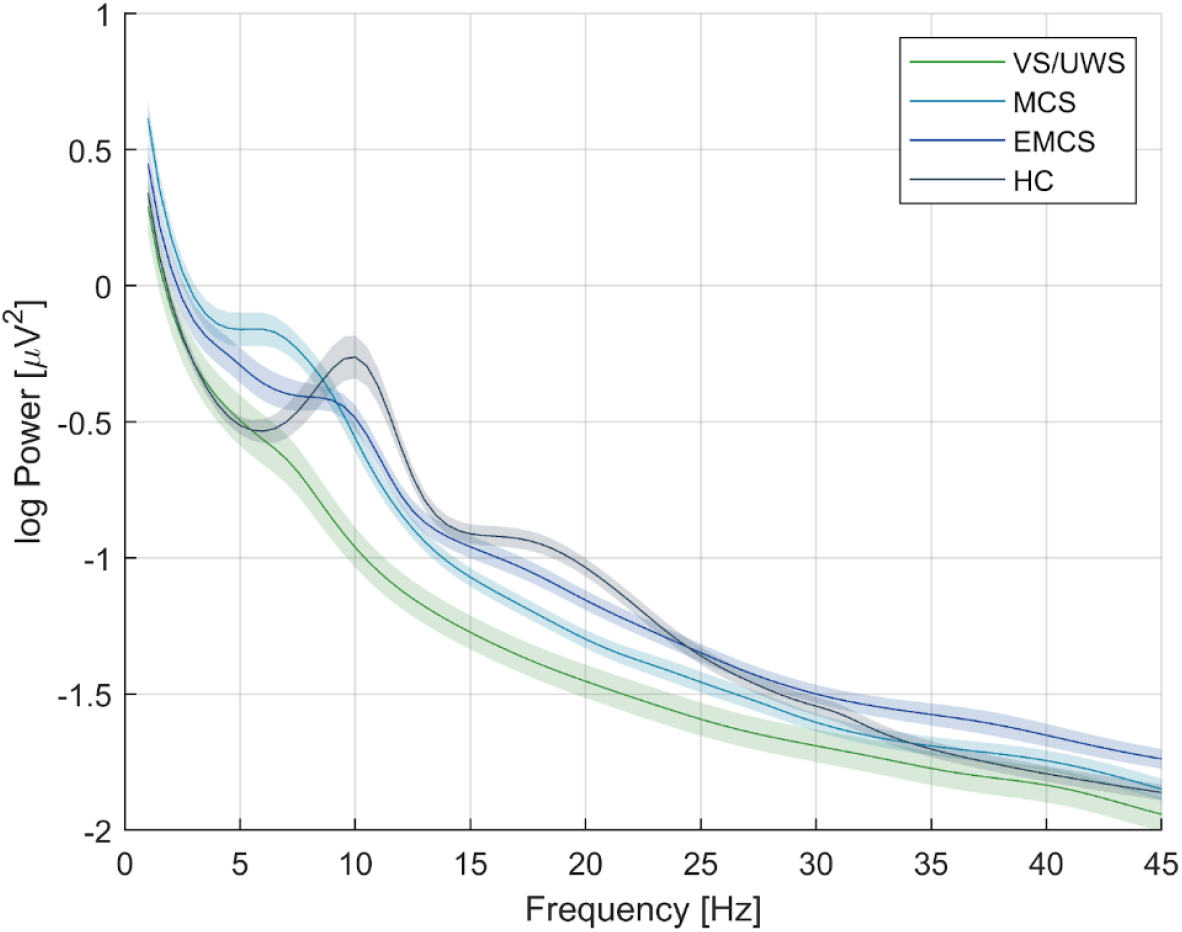
The averaged power spectra (*solid lines*) modeled with FOOOF parameters of all patient PDOC groups (UWS - unresponsive wakefulness syndrome, MCS - minimally conscious state, EMCS - emergence from MCS) and the healthy control group (HC). *Coloured ribbon* represents 95% SE around the fit.

**Figure 2.**
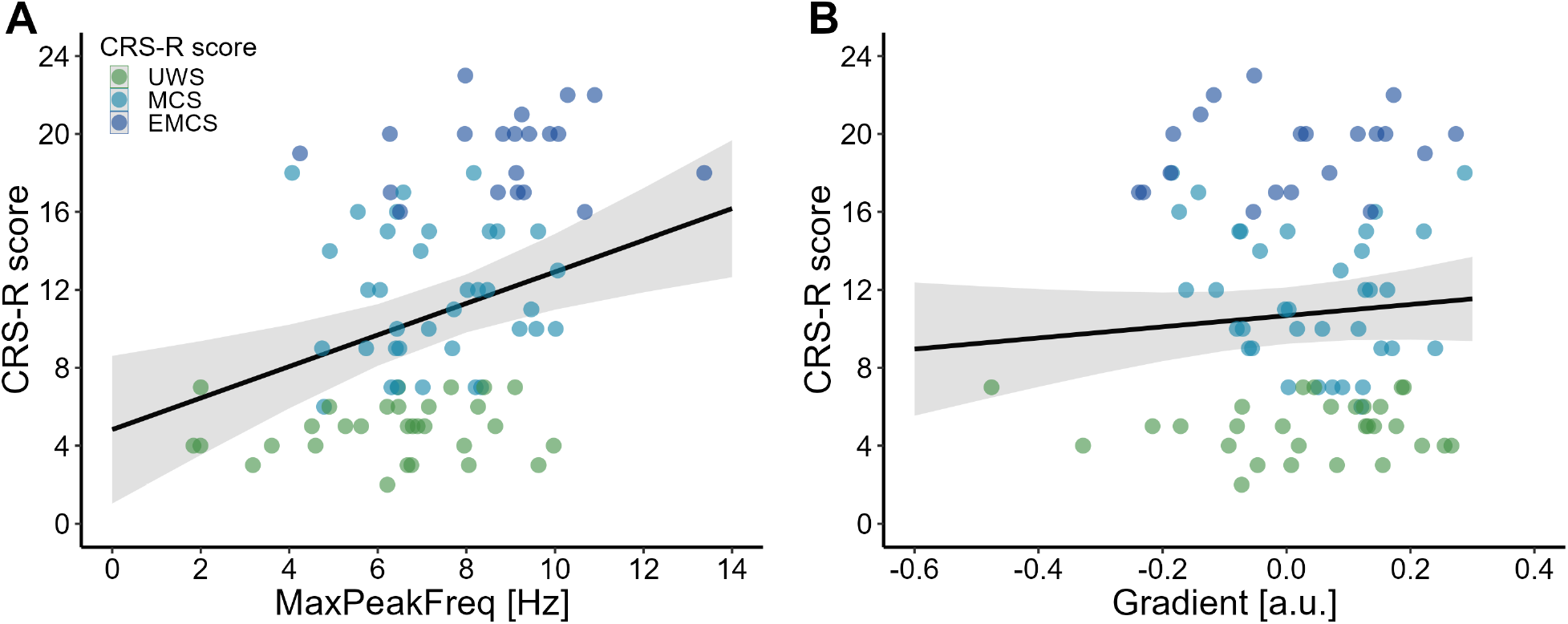
The relationship between CRS-R score value and **A** the frequency of the maximum peak in range 1-14 Hz (*MaxPeakFreq*) with the *Model 1a* fit, **B** antero-posterior gradient (*Gradient*) values with the *Model 1b* fit (*black solid line*), both performed on data from PDOC group. *Gray ribbon* represents 95% SE around the fit.

### Diagnostic utility of the AP gradient and the frequency of the maximum peak

The size of each of four groups included in the analysis, together with its demographic statistics is presented in Table **1**.

Among all fitted *CRSdiagnosis* models, *Model 2a* performed best compared to null model (Δ*LL* = −23.10, see Tab. **3**). Analysis showed a statistically significant effect of the highest peak in the 1-14 Hz range (*MaxPeakFreq* : *β* = 0.75 ± 0.14, *CI* : [0.48 1.03], Fig. **3**A). The probability of UWS classification decreases with the increasing *MaxPeakFreq* (*IQR* = 2.88) and approaches zero, with *MaxPeakFreq* value >8.5 Hz. Probability distribution of MCS assignment presents more Gaussian-like shape, with the maximal frequency of the highest peak around 7 Hz, (*IQR* = 2.08). For the *MaxPeakFreq* (*IQR* = 1.95) EMCS diagnosis probability increases only to the maximal point of 9 Hz, and starts decreasing. Finally, as predicted, HC profile presents an increase of *MaxPeakFreq* (*IQR* = 1.62) with the logarithmic increase of classification probability (see Fig. **3**B). *MaxPeakFreq* increased by 1 Hz multiplies the odds of HC diagnosis by 2.14. In the case of the AP gradient (*Model 2b*) and its interaction with the *MaxPeakFreq* (*Model 2c*) results did not reach significance threshold (*CI* : [-1.25 4.57] and *CI* : [-1.00 1.17], respectively, see Tab. **3**).

**Table 3.** Summary of the *CRSdiagnosis* models fitted with its accuracy diagnostic.

| Model | Effects | Estimate ± SE | z | p | 95% CI | $\sigma^2$ | $\tau_{00}$ | LL |
| --- | --- | --- | --- | --- | --- | --- | --- | --- |
| Model 2n | UWS \| MCS | -2.92 ± 0.74 | -3.95 | <0.001 | [-4.37 -1.47] | 1.00 | 12.14 | -148.59 |
| | MCS \| EMCS | -0.39 ± 0.50 | -0.79 | 0.432 | [-1.36 0.58] | | | |
| | EMCS \| HC | 1.13 ± 0.49 | 2.28 | 0.022 | [0.16 2.10] | | | |
| Model 2a | UWS \| MCS | 4.10 ± 0.93 | 4.39 | <0.001 | [2.27 5.94] | 1.00 | 2.84 | -125.49 |
| | MCS \| EMCS | 6.18 ± 1.16 | 5.33 | <0.001 | [3.90 8.45] | | | |
| | EMCS \| HC | 7.52 ± 1.38 | 5.44 | <0.001 | [4.81 10.22] | | | |
| | MaxPeakFreq | 0.75 ± 0.14 | 5.41 | <0.001 | [0.48 1.03] | | | |
| Model 2b | UWS \| MCS | -2.76 ± 0.71 | -3.85 | <0.001 | [-4.17 -1.36] | 1.00 | 10.86 | -147.99 |
| | MCS \| EMCS | -0.31 ± 0.47 | -0.66 | 0.510 | [-1.24 0.62] | | | |
| | EMCS \| HC | 1.15 ± 0.47 | 2.43 | 0.015 | [0.22 2.08] | | | |
| | Gradient | 1.66 ± 1.48 | 1.12 | 0.264 | [-1.25 4.57] | | | |
| Model 2c | UWS \| MCS | 4.23 ± 0.98 | 4.30 | <0.001 | [2.30 6.16] | 1.00 | 2.11 | -122.02 |
| | MCS \| EMCS | 6.18 ± 1.19 | 5.19 | <0.001 | [3.85 8.52] | | | |
| | EMCS \| HC | 7.47 ± 1.40 | 5.34 | <0.001 | [4.73 10.21] | | | |
| | MaxPeakFreq : Gradient | 0.09 ± 0.55 | 0.16 | 0.876 | [-1.00 1.17] | | | |

LL Log-Likelihood, $\sigma^2$ variance of level-1 residual errors, $\tau_{00}$ variance of level-2 residual errors.

**Figure 3.**
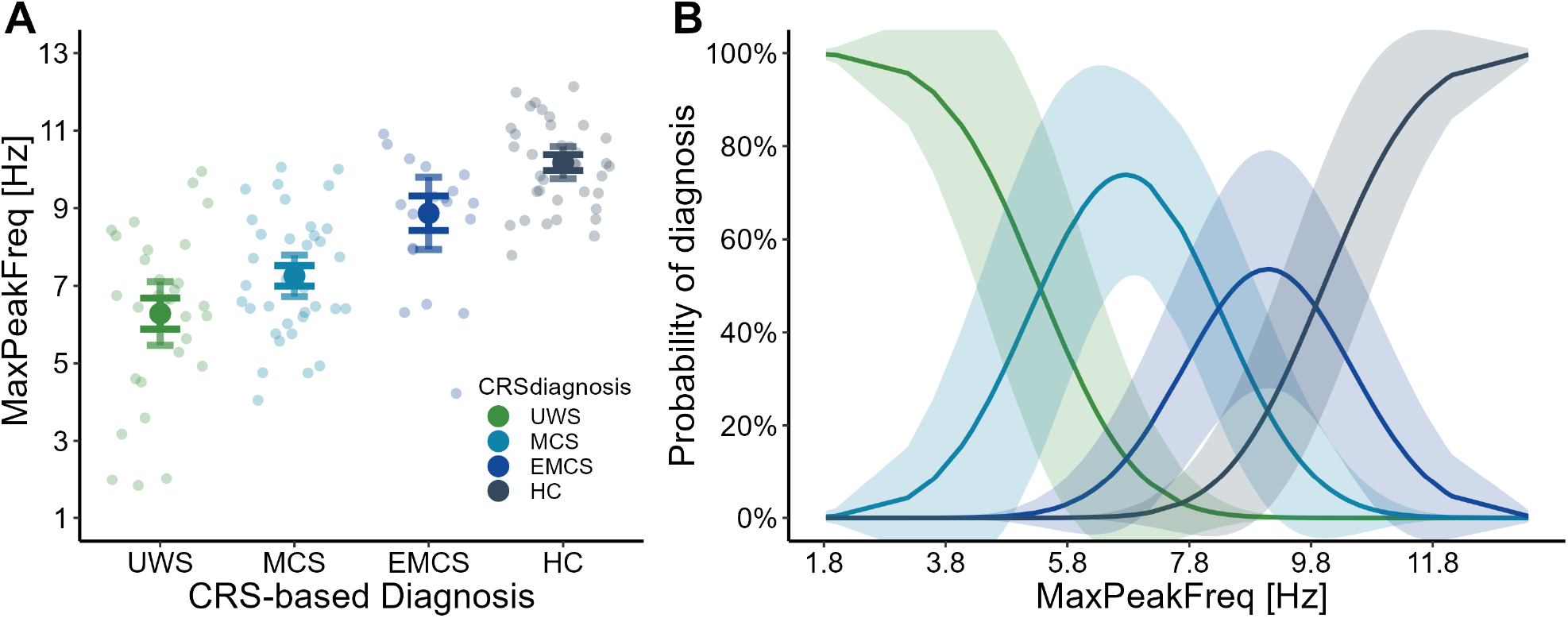
**A** The relationship between the frequency of the maximum peak in range 1-14 Hz (*MaxPeakFreq*) and CRS-based diagnosis including UWS (unresponsive wakefulness syndrome), MCS (minimally conscious state), EMCS (emergence from minimally conscious state) and HC (healthy control). *Light coloured dots* represent raw data points, *dark coloured dots* represent mean values for each diagnosis. Whiskers represent 95% SE (*dark coloured*) and 95% CE (*light coloured*) intervals. **B** Probability of classification to diagnosis groups (UWS, MCS, EMCS, HC) based on *MaxPeakFreq* as *solid lines*. *Coloured ribbon* represents 95% SE around the fit.

### The influence of the number of detected maxima on the postulated markers of neurocognitive recovery

The power spectrum analysis of the resting-state data revealed diversity in the number of peaks detected in the 1-14 Hz range. To account for the possible relation between the number of detected peaks and the predictive power of our factors, we divided the dataset into two groups. The first group presented a single maximal peak, while the second group included spectra with at least two peaks in the 1-14 Hz range (demographic statistics in Tab. B1 Appendix B of the Supplementary information). All previously used models for *CRSscore* (*Model 1n/a/b/c*) and *CRSdiagnosis* (*Model 2n/a/b/c*) were then refitted to both groups, separately (indicated as *s* prefix in the model’s name for the single peak data and as *m* prefix for the multiple peak data). The analysis revealed interesting differences between datasets. In the *CRSscore* models using a single-peak as well as multi-peak data, only the *MaxPeakFreq* factor reached a significance level (*p* = 0.003, see *Model s1a* in Tab. B2 and *p* = 0.010, see *Model m1a* in Tab. B3, respectively).

However, in the *CRSdiagnosis* single-peak model, both *MaxPeakFreq* and *Gradient* presented a diagnostic potential, with *p* = 0.017 (see *Model s2a* in Tab. B2) and *p* = 0.018 (see *Model s2b* in Tab. B2), respectively. For the multi-peak data, again only the *MaxPeakFreq* variable maintained a high significance level (*p* < 0.001, see *Model m2a* in Tab. B3). Dividing the dataset using the criterion of the number of maximal peaks in the searched EEG frequency range revealed a new aspect of the *Gradient* parameter. As demonstrated by the results for the patient spectra containing a single peak in the 1-14 Hz range, the antero-posterior gradient becomes a significant predictor of neurocognitive state (*Model s2b*). In such a case it seems that both the *Gradient* parameter and *MaxPeakFreq* are indicators of the same neurophysiological process.

To account for a possible influence of etiology and drug intake on our results we have conducted additional analyses with inclusion of etiology (anoxia, stroke, trauma). This analysis did not reveal any significant influence of that factor (see Appendix C of the Supplementary information for details). Furthermore, for a proportion of patients we were able to identify medications with possible influence on the central nervous system: Amantix (active substance *Amantadine Sulfate*), Baclofen (*Baclofen*) and Depakine (*Valproate*). An extended model which included those substances and patients without medications did not improve prediction capacity of any of the dependent variables (see Appendix D of the Supplementary information for details).

## Discussion

In this study, we have demonstrated that the frequency of the maximum peak in the EEG spectrum in the centro-parietal regions is a good indicator of neurocognitive capacity in PDOC patients. We have found that the increase of the dominant EEG oscillation frequency within the 1-14 Hz frequency band predicts the increase of the total CRS-R score and the heightened probability of acquiring a more favorable diagnosis. Consequently, the slowing of the oscillations in the 1-14 Hz range is related to the less favorable PDOC diagnosis and the impairment of neurocognitive capacity indexed by the total CRS-R score.

For decomposition of the EEG spectral profiles, we used the FOOOF algorithm, which represents EEG spectra as a combination of two types of parameters. The first parameter, the aperiodic component, corresponds to the non-oscillatory aspect of the EEG signal appearing as the 1/f distribution/function over signal frequency. The second parameter, the oscillatory part of the EEG signal is fitted by FOOOF using the series of Gaussian distributions with center frequencies representing the maximal amplitude of oscillation at a given frequency band. This way of decomposing the EEG signal in the frequency domain allows for a more flexible representation of the variability of the EEG spectrum in human populations. The FOOOF method is beneficial especially in comparison with the more dominant approach based on the analysis of mean power values calculated across traditional EEG spectral bands (eg. Lutkenhoff et al 2020) which is based on a priori delimitation of the borders between EEG frequency bands.

While we have demonstrated that the primary oscillatory activity within the 1-14 Hz band can predict the neurocognitive state of the PDOC patient, we have also found that this primary activity can be confounded by additional oscillations in the same range. This oscillatory activity may stem from the structural or functional cerebral lesions, which often manifest as slow oscillations within the theta or delta range (Galovic et al 2017). Nevertheless, in the case of the EEG spectral profile marked by a single maximal peak in the 1-14 Hz range, we observed a relationship between the strength of AP gradient and CRS-R diagnosis. This trend was absent when considering the whole set of data, as well as when only the data with multiple maximal peaks was analyzed. Thus the success of the automatically assessed AP gradient value as a diagnostic factor depends on the integrity of the leading frequency in the EEG spectrum. Our results suggested that the existence of the second, emergent maximal peak in range 1-14 Hz weakens the contribution of the AP gradient. However, the relationship between the gradient and the frequency of the *MaxPeakFreq* suggests that both these indicators point at the single coherent neurophysiological process, which could be interpreted in terms of thalamo-cortical network connectivity. Previous research implies (Forgacs et al 2014, 2017) that the presence of this kind of oscillation in the alpha range reflects preserved cell survival, so the observed slowing of the posterior oscillations is probably dependent on the inefficient facilitation of the thalamocortical connections by the neuromodulatory systems innervating the thalamus and the cortex. On the basis of the observed relationships between spectral patterns and behavioral data (see Fig. **1**), three types of spectral profiles can be identified, which correspond to a distinct modes of cerebral functioning in PDOC patients:

***Type I:*** the dominance of aperiodic EEG activity. For this type, most of the signal energy is accumulated within delta range with a dominant profile 1/f in the spectrum. This spectral profile was observed mostly in the UWS group and can be interpreted as evidence for the low excitability of the cerebral cortex (Rosanova et al 2012) and could be linked to the extensive functional and structural thalamocortical dysfunction. The dominance of delta-range activity has been previously reported in the studies of resting-state EEG in severe cases of brain lesions with UWS/VS diagnosis (eg. Lehembre et al 2012; Lutkenhoff et al 2020). However most of those studies were based on analyzing separate EEG spectral bands, so the 1/f pattern could be mistakenly treated as the increased oscillatory delta activity. The origins of aperiodic signals (1/f pattern) in human EEG are not well understood, however they are interpreted as the result of a shift in the balance between the synaptic excitation and inhibition (Donoghue et al 2020; Gao et al 2017).

***Type II:*** the single theta-alpha oscillation (4-14 Hz). This type is represented by a single oscillatory peak in the theta-alpha range within the centro-parietal region. Its frequency (the *MaxPeakFreq* parameter) is correlated with the PDOC diagnosis and total CRS-R score. Moreover, the gradient of the amplitude of this oscillation contributes to the prediction accuracy of the *MaxPeakFreq*. This pattern represents the strongest connection to the patient’s neurocognitive state as the frequency of oscillation, along with the prominence of AP gradient, most strongly predicts the PDOC diagnosis and CRS-R score in the group of patients where a single peak was detected within the 1-14 Hz range. Interestingly, it remained within 1-5 Hz range in UWS/VS patients, it ranged within the 5-8 Hz in MCS patients, and reached up to 8.5-10 Hz range in EMCS patients. In our opinion, despite its variable frequency, this oscillation represents a single neurophysiological process, related to the oscillation which, in normal awake EEG, is known as the posterior-dominant rhythm (PDR, Louis and Frey 2016). PDR slowing can be observed under several conditions, but its mechanisms are not fully understood (O’Gorman et al 2013). First, it is a normal developmental phenomenon that occurs during childhood (Marcuse et al 2016; Rodríguez-Martínez et al 2017; Cellier et al 2021). Second, it has been shown to accompany various conditions of neurological pathology, such as mild traumatic injury (Galovic et al 2017), white matter damage caused by vascular dementia (Moretti 2004) or decreased cerebral metabolism (Ingvar et al 1976; O’Gorman et al 2013). Finally it could be related to global brain pathology such as mild cognitive impairment or Alzheimer’s disease (Garcés et al 2013; Babiloni et al 2015; Benz et al 2014; Zimmermann et al 2015; Dickinson et al 2018).

***Type III:*** multiple theta-alpha oscillations (4-14 Hz). The characteristic feature of this type is a presence of multiple peaks within theta-alpha frequency range. This spectral pattern revealed weaker association with the patient’s neurocognitive state than Type II. We consider this profile as a mixture of type II oscillation and other slow oscillations. If present, these slow oscillations are probably caused by the structural damage to cortical mantle or the underlying subcortical structures. The presence of local slow rhythms in the vicinity of the lesion has been observed repeatedly in previous studies (e.g. Sarasso et al 2020). However, in the present study we do not have precise information about lesion location, so we cannot directly confirm this assumption. Note that the multiple oscillation pattern was not associated with the clinical status (CRS-R). This suggests that those additional oscillations are not related to the neural mechanisms influencing the clinical status, but are likely due to the more focal type of brain pathology. Moreover, in the case of type III the AP gradient data did not contribute to prediction of the state, confirming the presumption that they represent neural mechanisms which are of a different kind that those represented by the type II.

An attempt to investigate possible features of resting-state EEG activity on the basis of EEG spectral profiles has been previously made by Forgacs et al. (2014; 2017; Schiff et al 2014) in anoxic PDOC patients. These authors distinguished four categories of frequency spectrum profiles in PDOC patients (the ‘ABCD’ model) and linked those to the anoxic patients’ recovery. Crucially, in Forgacs et al. studies the spectral profiles were identified on the basis of qualitative experts evaluation. Their A-profile is very similar to our type I profile, with a dominance of 1/f power distribution across frequencies and association with UWS/VS diagnosis. The type B and D profiles identified by Forgacs et al. (2014) overlap with our type II profile, with the presence of a single peak within the theta-alpha band. In their studies these profiles most frequently appeared in MCS patients. The C-profile represented prominent peaks in theta in the posterior and beta activity in the frontal regions. Beta oscillations present in our sample were contaminated by frequent muscle artifacts in frontal channels. As we were not able to achieve the sufficient signal quality in the beta band, we could not corroborate Forgacs et al. (2014; 2017) findings within this frequency range.

In contrast with the distinctions proposed by Forgacs et al. (2014; 2017) we did not dissociate between spectral profiles with the single dominant theta and alpha activity, since our results suggest that they may represent a uniform neural mechanism in PDOC patients, based on the functional recovery of neuronal populations within the thalamocortical system. In this way, the process of regaining consciousness is based on the changes in dynamics of neural activity, resulting in the modulation of the frequency of the posterior dominant rhythm. The possible mechanism of this variability might involve dynamical changes in inhibition-excitation ratio within the thalamocortical system (Bhattacharya et al 2011). We support our conjecture with the observation that the dominant rhythm in the theta-alpha range (type II spectral profile) was accompanied with the presence of the anterior-posterior gradient despite the frequency range that was lower than the ‘canonical’ alpha range (namely 8-13 Hz).

Due to the numerous comorbidities, including epilepsy, parkinsonian tremor and muscle spasms, patients with disorders of consciousness are often treated with extensive pharmacological therapy. Some of those medications are reported to influence the EEG signal (Scarpino et al 2019; Mecarelli et al 2019; Ciurleo et al 2013; Cho et al 2012; Horiguchi et al 1990; Marciani et al 1993; Neckelmann et al 1996; Salinsky et al 2002). A particular pharmacotherapy is often linked to a specific etiology. Although etiology is not a direct indicator of a resulting disorder and a corresponding structural brain damage (Braakman et al 1988; Levin et al 1991; Levy et al 1981; Sazbon et al 1993; Sazbon and Groswasser 1990; Bates 1991; Shewmon and Giorgio 1989), it was shown to have some impact on the neurocognitive state (Forgacs et al 2020; Mecarelli et al 2019; Schiff 2010). To account for the possibility of relation between measured EEG characteristics and patient’s etiology or received medication, we compared the final models with its versions taking the effect of patients Etiology (anoxia, stroke and trauma) and chosen Medications (Amantix, Baclofen, Depakine, none) into account. None of the analyses revealed a robust significance of the Etiology or Medications groups in *CRSscore* and *CRSdiagnosis* models (see Appendix C and Appendix D in Supplementary information for details). This analysis ruled out etiology and drug intake as the significant interfering factors. Investigating individual power spectra based on extraction of frequency parameters within the 1-14 Hz range offers the possibility of complementing the diagnosis of the PDOC patient with a relatively simple tool. At the same time, however, the topographical patterns of dominant EEG oscillations should also be examined, as they could provide means of disentanglement between the EEG oscillations that have predictive power for patient state and other EEG oscillations that are also related to focal or global brain dysfunction, but did not show a clear relation to the diagnosis of PDOC.

## Conclusions

In this paper we investigated the possibility of improving the PDOC diagnostic process by introducing electrophysiological resting-state markers of patients’ neurocognitive capacity, based on the posterior dominant peak frequency and evaluation of the anterior-posterior gradient of posterior oscillations. Both parameters stand as a promising diagnostic tool based on the qualitative approach used in other resting state studies in the PDOC population. We have identified three types of spectral profiles, linked with different brain pathology. Profiles were closely associated with distinct PDOC diagnoses. FOOOF algorithm of spectral decomposition, which controls for the aperiodic signal background, allowed us to efficiently capture characteristics of frequency distribution and observed differences between spectra with single and multiple maximal peaks.

Peak frequency and gradient were calculated automatically from the EEG signal. Our work complements previously established PDOC profiles by including various etiology types as well as healthy control samples for the comparison. It presents a viable proposal of a simple yet efficient method of distinguishing between spectral PDOC profiles and may provide the means to explore its underlying mechanisms.

## Acknowledgments

This study was was supported by OPUS grants 2013/11/B/HS6/01242 and 2018/31/B/HS6/03920 from the National Science Center of Poland. The authors thank personnel of the patient rehabilitation centers in Torun, Czestochowa and Krakow for their invaluable help.

