## Supplementary information for "Distinct spectral profiles of awake resting EEG in disorders of consciousness - the role of frequency and topography of oscillations"

---

Dominika Drążyk [1], Karol Przewrocki [2], Urszula Górska [3], Marek Binder [4,*],

[1] *Institute of Neurosciences, Université catholique de Louvain, Brussels, Belgium*
[2] *Donders Institute for Brain, Cognition and Behavior, Radboud University Nijmegen, Netherlands*
[3] *Department of Psychiatry, University of Wisconsin-Madison, 53719 USA*
[4] *Institute of Psychology, Jagiellonian University, Krakow, Poland*

* corresponding author:
*Marek Binder, PhD,*
*Institute of Psychology, Ingardena 6, 30-060 Krakow, Poland*

### Supplementary information

#### Appendix A

##### Clinical and demographic characteristics of patients.

**Table A.1.** Clinical and demographic characteristics of patients.

| *Patient* | *Sex* | *Etiology* | *Session* | *Age* | *Time since injury [months]* | *CRS-R measurements date* | *EEG measurements date* | *AS* | *VS* | *MS* | *OS* | *CS* | *ARS* | *CRSscore* | *Diagnosis* | *Baclofen* | *Amantix* | *Depakine* | *Exclusion criteria* |
|---|---|---|---|---|---|---|---|---|---|---|---|---|---|---|---|---|---|---|---|
| P01 | M | anoxia | 1 | 18 | 38 | 2015-05-22 | 2015-05-25 | 2 | 0 | 2 | 0 | 0 | 2 | 6 | UWS | N/A | N/A | N/A | artifacts in the midline data |
| P02 | M | anoxia | 1 | 66 | 18 | 2015-05-11 | 2015-05-27 | 2 | 4 | 5 | 2 | 1 | 2 | 16 | MCS | N/A | N/A | N/A | |
| | | | 2 | 66 | 23 | 2015-10-09 | 2015-09-29 | 3 | 5 | 2 | 2 | 1 | 2 | 15 | MCS | 0 | 0 | 0 | |
| | | | 3 | 67 | 30 | 2016-05-05 | 2016-05-10 | 3 | 5 | 2 | 2 | 1 | 2 | 15 | MCS | N/A | N/A | N/A | artifacts in the midline data |
| | | | 4 | 67 | 31 | 2016-06-06 | 2016-06-07 | 3 | 5 | 2 | 2 | 1 | 2 | 15 | MCS | N/A | N/A | N/A | extensive artifacts in the data |
| | | | 5 | 68 | 39 | 2017-01-31 | 2017-01-31 | 2 | 3 | 5 | 2 | 0 | 3 | 15 | MCS | 0 | 0 | 0 | extensive artifacts in the data |
| P03 | F | N/A | 1 | 43 | 8 | 2015-05-20 | 2015-05-26 | 2 | 3 | 2 | 2 | 1 | 2 | 12 | MCS | N/A | N/A | N/A | |
| | | | 2 | 44 | 16 | 2016-01-21 | 2016-01-27 | 2 | 3 | 2 | 1 | 0 | 1 | 9 | MCS | 1 | 0 | 0 | |
| P04 | M | trauma | 1 | 65 | 12 | 2015-05-20 | 2015-05-26 | 1 | 3 | 2 | 2 | 2 | 2 | 12 | MCS | N/A | N/A | N/A | |
| | | | 2 | 65 | 16 | 2015-10-06 | 2015-09-29 | 1 | 2 | 1 | 1 | 0 | 1 | 6 | UWS | N/A | N/A | N/A | artifacts in the midline data |
| P05 | M | trauma | 1 | 58 | 8 | 2015-05-20 | 2015-05-27 | 2 | 0 | 0 | 0 | 0 | 2 | 4 | UWS | N/A | N/A | N/A | |
| P06 | F | trauma | 1 | 51 | 18 | 2015-06-08 | 2015-05-28 | 4 | 5 | 5 | 2 | 2 | 3 | 21 | EMCS | N/A | N/A | N/A | |
| P07 | M | anoxia | 1 | 26 | 17 | 2015-10-08 | 2015-09-30 | 1 | 0 | 1 | 0 | 0 | 2 | 4 | UWS | N/A | N/A | N/A | |
| P08 | M | trauma | 1 | 31 | 6 | 2015-06-08 | 2015-05-29 | 3 | 3 | 5 | 2 | 2 | 3 | 18 | MCS | 1 | 0 | 0 | |
| | | | 2 | 31 | 10 | 2015-10-07 | 2015-09-30 | 3 | 5 | 2 | 1 | 1 | 3 | 15 | MCS | 1 | 0 | 0 | |
| P09 | F | trauma | 1 | 21 | 16 | 2015-05-22 | 2015-05-29 | 4 | 5 | 5 | 2 | 1 | 1 | 18 | MCS | N/A | N/A | N/A | artifacts in the midline data |
| P10 | M | anoxia | 1 | 37 | 22 | 2015-07-03 | 2015-06-24 | 1 | 0 | 1 | 0 | 0 | 2 | 4 | UWS | N/A | N/A | N/A | artifacts in the midline data |
| | | | 2 | 38 | 29 | 2016-02-05 | 2016-02-05 | 2 | 1 | 2 | 0 | 0 | 2 | 7 | UWS | 1 | 0 | 1 | |
| P11 | M | anoxia | 1 | 48 | 108 | 2016-01-19 | 2016-01-25 | 1 | 1 | 0 | 0 | 0 | 1 | 3 | UWS | N/A | N/A | N/A | artifacts in the midline data |
| P12 | M | anoxia | 1 | 51 | 67 | 2015-07-03 | 2015-06-25 | 1 | 0 | 1 | 1 | 0 | 1 | 4 | UWS | N/A | N/A | N/A | artifacts in the midline data |
| | | | 2 | 52 | 76 | 2016-01-20 | 2016-01-26 | 1 | 0 | 1 | 0 | 0 | 1 | 3 | UWS | 1 | 0 | 0 | |
| P13 | M | anoxia | 1 | 39 | 50 | 2015-07-03 | 2015-06-26 | 1 | 0 | 2 | 1 | 0 | 1 | 5 | UWS | N/A | N/A | N/A | |
| | | | 2 | 40 | 57 | 2016-01-22 | 2016-01-28 | 1 | 0 | 2 | 0 | 0 | 1 | 4 | UWS | 0 | 0 | 0 | |
| P14 | M | anoxia | 1 | 59 | 8 | 2016-01-19 | 2016-01-25 | 1 | 0 | 1 | 0 | 0 | 1 | 3 | UWS | 1 | 0 | 0 | |
| P15 | M | anoxia | 1 | 27 | 67 | 2015-07-01 | 2015-06-26 | 3 | 3 | 2 | 2 | 1 | 3 | 14 | MCS | N/A | N/A | N/A | |
| | | | 2 | 28 | 80 | 2016-01-04 | 2016-01-27 | 4 | 5 | 5 | 1 | 2 | 3 | 20 | EMCS | 1 | 0 | 0 | |
| | | | 3 | 28 | 86 | 2016-06-06 | 2016-06-06 | 4 | 5 | 6 | 2 | 2 | 3 | 22 | EMCS | N/A | N/A | N/A | |
| P16 | M | anoxia | 1 | 51 | 17 | 2015-10-09 | 2015-10-01 | 1 | 0 | 0 | 0 | 0 | 2 | 3 | UWS | N/A | N/A | N/A | artifacts in the midline data |
| | | | 2 | 52 | 24 | 2016-05-10 | 2016-05-09 | 1 | 0 | 1 | 0 | 0 | 1 | 3 | UWS | N/A | N/A | N/A | artifacts in the midline data |
| | | | 3 | 52 | 25 | 2016-06-07 | 2016-06-07 | 1 | 0 | 1 | 0 | 0 | 1 | 3 | UWS | N/A | N/A | N/A | artifacts in the midline data |
| | | | 4 | 53 | 32 | 2017-01-31 | 2017-01-31 | 1 | 0 | 1 | 0 | 0 | 1 | 3 | UWS | 0 | 0 | 0 | |
| P17 | F | trauma | 1 | 24 | 15 | 2015-10-09 | 2015-10-02 | 1 | 0 | 2 | 1 | 0 | 1 | 5 | UWS | 0 | 0 | 1 | extensive artifacts in the data |
| | F | | 2 | 25 | 22 | 2016-05-09 | 2016-05-10 | 1 | 0 | 1 | 1 | 0 | 1 | 4 | UWS | N/A | N/A | N/A | |
| | F | | 3 | 25 | 23 | 2016-06-08 | 2016-06-08 | 1 | 0 | 1 | 1 | 0 | 1 | 4 | UWS | N/A | N/A | N/A | |
| P18 | M | N/A | 1 | 28 | 6 | 2016-02-05 | 2016-01-29 | 1 | 0 | 1 | 0 | 0 | 1 | 3 | UWS | 1 | 0 | 1 | artifacts in the midline data |
| | | | 2 | 28 | 11 | 2016-06-07 | 2016-06-07 | 1 | 0 | 1 | 0 | 0 | 0 | 2 | UWS | N/A | N/A | N/A | artifacts in the midline data |
| P19 | F | stroke | 1 | 37 | 5 | 2016-02-05 | 2016-01-29 | 4 | 5 | 4 | 1 | 2 | 2 | 18 | EMCS | 0 | 1 | 0 | artifacts in the midline data |
| | | | 2 | 37 | 13 | 2016-10-03 | 2016-10-03 | 4 | 5 | 2 | 1 | 2 | 3 | 17 | EMCS | N/A | N/A | N/A | |
| | | | 3 | 38 | 18 | 2017-02-01 | 2017-02-01 | 4 | 5 | 2 | 1 | 2 | 3 | 17 | EMCS | 0 | 0 | 1 | |
| P20 | M | trauma | 1 | 37 | 11 | 2016-05-06 | 2016-05-11 | 1 | 0 | 1 | 0 | 0 | 0 | 2 | UWS | N/A | N/A | N/A | |
| | | | 2 | 37 | 16 | 2016-10-05 | 2016-10-05 | 1 | 0 | 1 | 0 | 0 | 1 | 3 | UWS | N/A | N/A | N/A | |
| P21 | M | trauma | 1 | 21 | 8 | 2016-05-16 | 2016-05-11 | 4 | 5 | 2 | 2 | 1 | 1 | 15 | MCS | N/A | N/A | N/A | |
| P22 | M | trauma | 1 | 19 | 6 | 2016-05-09 | 2016-05-12 | 4 | 5 | 1 | 2 | 2 | 2 | 16 | MCS | N/A | N/A | N/A | |
| P23 | M | N/A | 1 | 29 | 6 | 2016-05-17 | 2016-05-12 | 1 | 0 | 2 | 0 | 0 | 2 | 5 | UWS | N/A | N/A | N/A | |
| | | | 2 | 29 | 11 | 2016-10-05 | 2016-10-05 | 1 | 0 | 3 | 1 | 0 | 2 | 7 | MCS | N/A | N/A | N/A | |
| | | | 3 | 30 | 15 | 2017-01-30 | 2017-01-30 | 1 | 3 | 3 | 1 | 0 | 2 | 10 | MCS | 1 | 0 | 1 | |
| | | | 4 | 30 | 23 | 2017-10-05 | 2017-10-05 | 1 | 1 | 1 | 1 | 0 | 1 | 5 | UWS | N/A | N/A | N/A | |
| P24 | F | trauma | 1 | 25 | 6 | 2016-05-13 | 2016-05-13 | 1 | 1 | 1 | 0 | 0 | 2 | 5 | UWS | N/A | N/A | N/A | |
| | | | 2 | 26 | 19 | 2017-10-02 | 2017-10-02 | 1 | 1 | 2 | 1 | 0 | 2 | 7 | UWS | N/A | N/A | N/A | |
| | | | 3 | 26 | 23 | 2019-08-09 | 2019-08-09 | 1 | 0 | 1 | 0 | 0 | 1 | 3 | UWS | N/A | N/A | N/A | |
| | | | 4 | 28 | 45 | 2019-10-25 | 2019-10-25 | 3 | 1 | 2 | 1 | 0 | 2 | 9 | MCS | 0 | 0 | 0 | |
| | | | 5 | 28 | 46 | 2019-12-23 | 2019-12-23 | 1 | 3 | 2 | 1 | 1 | 2 | 10 | MCS | 0 | 0 | 0 | |
| | | | 6 | 28 | 48 | 2019-10-25 | 2019-10-25 | 1 | 1 | 2 | 1 | 0 | 2 | 7 | UWS | N/A | N/A | N/A | artifacts in the midline data |
| | | | 7 | 28 | 50 | 2019-12-23 | 2019-12-23 | 3 | 1 | 2 | 1 | 0 | 2 | 9 | MCS | N/A | N/A | N/A | |
| P25 | M | anoxia | 1 | 37 | 10 | 2016-05-12 | 2016-05-13 | 2 | 0 | 1 | 0 | 0 | 2 | 5 | UWS | N/A | N/A | N/A | extensive artifacts in the data |
| | | | 2 | 37 | 15 | 2016-10-03 | 2016-10-03 | 1 | 0 | 1 | 0 | 0 | 2 | 4 | UWS | N/A | N/A | N/A | artifacts in the midline data |
| P26 | F | stroke | 1 | 38 | 8 | 2016-10-04 | 2016-10-04 | 4 | 5 | 1 | 1 | 2 | 3 | 16 | EMCS | N/A | N/A | N/A | |
| P27 | M | trauma | 1 | 55 | 14 | 2016-06-07 | 2016-06-08 | 4 | 5 | 4 | 1 | 2 | 3 | 19 | EMCS | N/A | N/A | N/A | |
| | | | 2 | 56 | 22 | 2017-01-31 | 2017-01-30 | 4 | 5 | 4 | 1 | 1 | 3 | 18 | MCS | 0 | 0 | 0 | |
| P28 | F | trauma | 1 | 30 | 8 | 2016-06-07 | 2016-06-08 | 1 | 1 | 1 | 1 | 0 | 2 | 6 | UWS | N/A | N/A | N/A | |
| P29 | M | anoxia | 1 | 33 | 19 | 2016-10-04 | 2016-10-04 | 1 | 0 | 1 | 0 | 0 | 1 | 3 | UWS | N/A | N/A | N/A | artifacts in the midline data |
| | | | 2 | 34 | 23 | 2017-02-02 | 2017-02-02 | 1 | 0 | 1 | 0 | 0 | 1 | 3 | UWS | 1 | 1 | 0 | extensive artifacts in the data |
| | | | 3 | 34 | 23 | 2017-02-02 | 2017-02-02 | 1 | 0 | 1 | 0 | 0 | 1 | 3 | UWS | 1 | 0 | 1 | extensive artifacts in the data |
| P30 | M | trauma | 1 | 65 | 3 | 2016-10-17 | 2016-10-17 | 3 | 4 | 2 | 1 | 1 | 1 | 12 | MCS | 1 | 1 | 0 | |
| P31 | M | trauma | 1 | 65 | 32 | 2016-10-17 | 2016-10-17 | 0 | 1 | 2 | 2 | 0 | 2 | 7 | UWS | N/A | N/A | N/A | artifacts in the midline data |
| | | | 2 | 66 | 40 | 2017-05-22 | 2017-05-22 | 1 | 0 | 2 | 2 | 0 | 2 | 7 | UWS | N/A | N/A | N/A | |
| P32 | M | anoxia | 1 | 47 | 6 | 2016-10-24 | 2016-10-24 | 3 | 0 | 1 | 1 | 1 | 1 | 7 | MCS | 0 | 1 | 0 | artifacts in the midline data |

| *Patient* | *Sex* | *Etiology* | *Session* | *Age* | *Time since injury [months]* | *CRS-R measurements date* | *EEG measurements date* | *AS* | *VS* | *MS* | *OS* | *CS* | *ARS* | *CRSscore* | *Diagnosis* | *Baclofen* | *Amantix* | *Depakine* | *Exclusion criteria* |
|---|---|---|---|---|---|---|---|---|---|---|---|---|---|---|---|---|---|---|---|
| P33 | F | anoxia | 1 | 23 | 2 | 2016-10-24 | 2016-10-24 | 3 | 1 | 2 | 1 | 1 | 3 | 11 | MCS | N/A | N/A | N/A | extensive artifacts in the data |
| P34 | F | anoxia | 1 | 18 | 2 | 2016-11-26 | 2016-11-26 | 1 | 0 | 2 | 1 | 0 | 1 | 5 | UWS | 1 | 0 | 0 | artifacts in the midline data |
| P35 | M | anoxia | 1 | 80 | 4 | 2016-11-26 | 2016-11-26 | 1 | 0 | 1 | 1 | 0 | 1 | 4 | UWS | 0 | 0 | 0 | artifacts in the midline data |
| | | | 2 | 81 | 9 | 2017-05-22 | 2017-05-22 | 1 | 0 | 1 | 1 | 0 | 1 | 4 | UWS | N/A | N/A | N/A | extensive artifacts in the data |
| P36 | F | trauma | 1 | 52 | 2 | 2016-12-05 | 2016-12-05 | 3 | 1 | 5 | 1 | 0 | 2 | 12 | MCS | 0 | 0 | 0 | |
| P37 | M | trauma | 1 | 18 | 36 | 2017-01-16 | 2017-01-16 | 1 | 1 | 2 | 1 | 0 | 1 | 6 | UWS | 0 | 0 | 1 | |
| P38 | M | anoxia | 1 | 37 | 8 | 2017-01-23 | 2017-01-23 | 1 | 0 | 2 | 1 | 0 | 2 | 6 | UWS | N/A | N/A | N/A | extensive artifacts in the data |
| P39 | M | anoxia | 1 | 11 | 21 | 2017-01-23 | 2017-01-23 | 2 | 2 | 5 | 2 | 0 | 2 | 13 | MCS | N/A | N/A | N/A | extensive artifacts in the data |
| P40 | F | anoxia | 1 | 62 | 2 | 2017-05-30 | 2017-05-30 | 1 | 0 | 2 | 1 | 0 | 1 | 5 | UWS | N/A | N/A | N/A | artifacts in the midline data |
| | | | 2 | 62 | 3 | 2017-10-04 | 2017-10-04 | 1 | 0 | 2 | 1 | 0 | 1 | 5 | UWS | N/A | N/A | N/A | extensive artifacts in the data |
| P41 | F | trauma | 1 | 43 | 1 | 2017-02-03 | 2017-02-03 | 1 | 1 | 1 | 0 | 0 | 1 | 4 | UWS | N/A | N/A | N/A | artifacts in the midline data |
| | | | 2 | 43 | 2 | 2017-06-01 | 2017-06-01 | 1 | 1 | 2 | 1 | 0 | 1 | 6 | UWS | N/A | N/A | N/A | |
| P42 | F | stroke | 1 | 49 | 1 | 2017-02-21 | 2017-02-21 | 1 | 3 | 2 | 1 | 1 | 1 | 9 | MCS | N/A | N/A | N/A | |
| P43 | M | trauma | 1 | 21 | 1 | 2017-02-21 | 2017-02-21 | 3 | 5 | 2 | 1 | 2 | 3 | 16 | EMCS | N/A | N/A | N/A | |
| | | | 2 | 21 | 2 | 2017-05-29 | 2017-05-29 | 3 | 3 | 5 | 1 | 2 | 3 | 17 | EMCS | N/A | N/A | N/A | |
| | | | 3 | 21 | 3 | 2017-11-08 | 2017-11-08 | 4 | 5 | 6 | 1 | 1 | 3 | 20 | EMCS | 0 | 0 | 0 | |
| P44 | F | trauma | 1 | 31 | 1 | 2017-02-22 | 2017-02-22 | 3 | 5 | 2 | 1 | 1 | 2 | 14 | MCS | N/A | N/A | N/A | |
| | | | 2 | 31 | 2 | 2017-05-30 | 2017-05-30 | 2 | 3 | 1 | 1 | 0 | 2 | 9 | MCS | N/A | N/A | N/A | |
| P45 | F | stroke | 1 | 75 | 1 | 2017-02-22 | 2017-02-22 | 2 | 3 | 5 | 3 | 1 | 1 | 15 | MCS | N/A | N/A | N/A | |
| P46 | M | trauma | 1 | 20 | 1 | 2017-03-27 | 2017-03-27 | 0 | 0 | 2 | 1 | 0 | 1 | 4 | UWS | N/A | N/A | N/A | artifacts in the midline data |
| P47 | F | trauma | 1 | 23 | 1 | 2017-03-27 | 2017-03-27 | 1 | 2 | 2 | 1 | 0 | 1 | 7 | MCS | N/A | N/A | N/A | |
| P48 | F | stroke | 1 | 31 | 1 | 2017-04-10 | 2017-04-10 | 3 | 1 | 5 | 2 | 1 | 3 | 15 | MCS | N/A | N/A | N/A | |
| | | | 2 | 32 | 2 | 2018-03-16 | 2018-03-15 | 3 | 4 | 6 | 2 | 1 | 2 | 18 | EMCS | 0 | 0 | 0 | |
| | | | 3 | 32 | 3 | 2018-03-16 | 2018-03-16 | 3 | 4 | 6 | 2 | 1 | 2 | 18 | EMCS | N/A | N/A | N/A | |
| P49 | M | anoxia | 1 | 34 | 1 | 2017-04-10 | 2017-04-10 | 1 | 1 | 1 | 1 | 0 | 2 | 6 | UWS | N/A | N/A | N/A | artifacts in the midline data |
| P50 | M | trauma | 1 | 25 | 1 | 2017-05-30 | 2017-05-29 | 2 | 3 | 2 | 1 | 0 | 2 | 10 | MCS | N/A | N/A | N/A | |
| | | | 2 | 25 | 2 | 2017-10-03 | 2017-10-03 | 2 | 3 | 2 | 2 | 0 | 2 | 11 | MCS | 1 | 0 | 0 | |
| | | | 3 | 25 | 3 | 2017-11-07 | 2017-11-07 | 2 | 3 | 2 | 1 | 0 | 2 | 10 | MCS | 1 | 0 | 0 | |
| P51 | M | trauma | 1 | 22 | 1 | 2017-05-29 | 2017-05-29 | 1 | 1 | 1 | 1 | 0 | 1 | 5 | UWS | N/A | N/A | N/A | |
| | | | 2 | 23 | 2 | 2018-03-12 | 2018-03-12 | 1 | 1 | 2 | 1 | 0 | 1 | 6 | UWS | 1 | 0 | 1 | |
| P52 | M | trauma | 1 | 24 | 1 | 2017-05-30 | 2017-05-30 | 4 | 5 | 6 | 3 | 1 | 3 | 22 | EMCS | N/A | N/A | N/A | |
| P53 | F | trauma | 1 | 37 | 1 | 2017-05-31 | 2017-05-31 | 3 | 3 | 1 | 1 | 0 | 3 | 11 | MCS | 1 | 1 | 0 | extensive artifacts in the data |
| | | | 2 | 37 | 2 | 2017-11-08 | 2017-11-08 | 4 | 5 | 2 | 1 | 2 | 3 | 17 | EMCS | 1 | 1 | 0 | |
| | | | 3 | 38 | 3 | 2018-03-12 | 2018-03-12 | 3 | 3 | 2 | 2 | 0 | 2 | 12 | MCS | 1 | 1 | 0 | |
| P54 | M | trauma | 1 | 34 | 1 | 2017-06-01 | 2017-06-01 | 1 | 0 | 1 | 1 | 0 | 2 | 5 | UWS | N/A | N/A | N/A | artifacts in the midline data |
| P55 | F | trauma | 1 | 35 | 1 | 2017-06-01 | 2017-06-01 | 1 | 0 | 1 | 1 | 0 | 2 | 5 | UWS | N/A | N/A | N/A | artifacts in the midline data |
| P56 | F | N/A | 1 | 23 | 1 | 2017-06-19 | 2017-06-19 | 1 | 1 | 1 | 2 | 0 | 2 | 7 | UWS | N/A | N/A | N/A | artifacts in the midline data |
| P57 | M | trauma | 1 | 30 | 1 | 2017-06-19 | 2017-06-19 | 1 | 1 | 2 | 1 | 0 | 2 | 7 | UWS | N/A | N/A | N/A | |
| P58 | F | trauma | 1 | 32 | 1 | 2017-10-02 | 2017-10-02 | 4 | 5 | 2 | 2 | 1 | 3 | 17 | MCS | 1 | 1 | 0 | |
| | | | 2 | 32 | 2 | 2017-11-08 | 2017-11-08 | 4 | 5 | 6 | 3 | 2 | 3 | 23 | EMCS | 1 | 1 | 1 | |
| P59 | M | trauma | 1 | 30 | 1 | 2017-10-02 | 2017-10-02 | 3 | 3 | 2 | 1 | 0 | 1 | 10 | MCS | 0 | 1 | 0 | |
| | | | 2 | 30 | 2 | 2017-11-07 | 2017-11-07 | 3 | 5 | 2 | 1 | 1 | 1 | 13 | MCS | 0 | 1 | 0 | |
| P60 | M | trauma | 1 | 26 | 1 | 2017-10-03 | 2017-10-03 | 4 | 5 | 5 | 1 | 2 | 3 | 20 | EMCS | 0 | 0 | 0 | |
| P61 | F | trauma | 1 | 22 | 1 | 2017-10-04 | 2017-10-04 | 2 | 3 | 2 | 1 | 0 | 2 | 10 | MCS | N/A | N/A | N/A | extensive artifacts in the data |
| | | | 2 | 22 | 2 | 2017-11-07 | 2017-11-07 | 2 | 3 | 2 | 1 | 0 | 2 | 10 | MCS | N/A | N/A | N/A | extensive artifacts in the data |
| P62 | M | anoxia | 1 | 28 | 1 | 2017-10-05 | 2017-10-05 | 1 | 1 | 2 | 1 | 0 | 2 | 7 | UWS | N/A | N/A | N/A | artifacts in the midline data |
| P63 | M | anoxia | 1 | 33 | 1 | 2017-10-05 | 2017-10-05 | 1 | 0 | 1 | 0 | 0 | 2 | 4 | UWS | N/A | N/A | N/A | artifacts in the midline data |
| P64 | M | trauma | 1 | 55 | 1 | 2018-02-09 | 2018-02-09 | 0 | 3 | 2 | 0 | 0 | 1 | 6 | MCS | 1 | 0 | 0 | |
| P65 | M | trauma | 1 | 30 | 1 | 2018-03-12 | 2018-03-12 | 1 | 0 | 2 | 2 | 0 | 1 | 6 | UWS | 0 | 0 | 0 | |
| P66 | F | stroke | 1 | 63 | 1 | 2018-03-13 | 2018-03-12 | 2 | 1 | 2 | 2 | 0 | 1 | 8 | UWS | N/A | N/A | N/A | extensive artifacts in the data |
| P67 | F | anoxia | 1 | 37 | 1 | 2018-03-13 | 2018-03-12 | 1 | 4 | 6 | 2 | 0 | 3 | 16 | EMCS | 1 | 0 | 1 | extensive artifacts in the data |
| P68 | M | trauma | 1 | 38 | 1 | 2018-03-16 | 2018-03-16 | 1 | 0 | 2 | 1 | 0 | 1 | 5 | UWS | 1 | 0 | 1 | |
| P69 | F | stroke | 1 | 32 | 1 | 2018-04-17 | 2018-04-17 | 2 | 1 | 2 | 1 | 0 | 1 | 7 | UWS | 0 | 0 | 1 | |
| P70 | M | anoxia | 1 | 59 | 1 | 2018-04-18 | 2018-04-27 | 1 | 0 | 0 | 2 | 0 | 2 | 5 | UWS | 0 | 0 | 0 | artifacts in the midline data |
| P71 | F | anoxia | 1 | 56 | 1 | 2018-09-11 | 2018-09-11 | 1 | 0 | 1 | 0 | 0 | 1 | 3 | UWS | N/A | N/A | N/A | extensive artifacts in the data |
| P72 | F | stroke | 1 | N/A | 1 | 2018-11-28 | 2018-11-28 | 3 | 0 | 5 | 1 | 1 | 1 | 11 | MCS | 0 | 0 | 0 | |
| P73 | M | trauma | 1 | 41 | 1 | 2018-11-30 | 2018-11-30 | 3 | 0 | 2 | 1 | 0 | 1 | 7 | MCS | 1 | 0 | 0 | |
| P74 | M | stroke | 1 | 55 | 1 | 2018-11-30 | 2018-11-30 | 1 | 0 | 2 | 0 | 0 | 2 | 5 | UWS | 1 | 0 | 1 | |
| P75 | M | anoxia | 1 | 39 | 2 | 2019-08-09 | 2019-08-09 | 1 | 0 | 2 | 0 | 0 | 1 | 4 | UWS | 0 | 0 | 0 | |
| | | | 2 | 39 | 3 | 2019-09-17 | 2019-09-17 | 1 | 0 | 2 | 1 | 0 | 1 | 5 | UWS | 0 | 0 | 0 | |
| P76 | F | anoxia | 1 | 41 | 1 | 2019-09-04 | 2019-09-04 | 1 | 0 | 1 | 1 | 1 | 2 | 6 | UWS | N/A | N/A | N/A | artifacts in the midline data |
| P77 | M | trauma | 1 | N/A | 1 | 2019-11-04 | 2019-11-04 | 4 | 5 | 5 | 0 | 3 | 3 | 20 | EMCS | 0 | 0 | 0 | |
| | | | 2 | N/A | 2 | 2019-12-03 | 2019-12-03 | 4 | 5 | 6 | 0 | 2 | 3 | 20 | EMCS | 0 | 0 | 0 | |
| P78 | F | stroke | 1 | 53 | 1 | 2019-11-04 | 2019-11-04 | 1 | 2 | 2 | 0 | 1 | 3 | 7 | MCS | 0 | 0 | 0 | |
| | | | 2 | 53 | 2 | 2019-12-03 | 2019-12-03 | 1 | 1 | 3 | 0 | 1 | 3 | 7 | MCS | 0 | 0 | 0 | |
| P79 | F | trauma | 1 | N/A | 1 | 2019-11-04 | 2019-11-04 | 4 | 5 | 6 | 0 | 2 | 3 | 20 | EMCS | 0 | 0 | 0 | |
| | | | 2 | N/A | 2 | 2019-12-03 | 2019-12-03 | 4 | 5 | 6 | 0 | 3 | 2 | 20 | EMCS | 0 | 0 | 0 | |

### Appendix B

**The influence of the number of detected maxima on the postulated markers of neurocognitive recovery.**

**Table B.1.** Demographics of patient groups within numbers of maximal peaks in the 1-14 Hz range.

| Number of maximal peaks in 1-14 Hz range | | | | | | |
|---|---|---|---|---|---|---|
| | *Groups* | *N measurements* | *N* | *Females* | *Age* | |
| | | | | | *M* | *SD* |
| Model 1 | single peak | 37 | 30 | 12 | 37.12 | 13.11 |
| | multiple peaks | 49 | 33 | 11 | 33.62 | 13.28 |
| Model 2 | single peak | 58 | 51 | 24 | 32.67 | 12.62 |
| | multiple peaks | 65 | 49 | 21 | 36.71 | 13.94 |

**Table B.2.** Results describing *CRSscore* and *CRSdiagnosis* models in a group of patients with single maximal peak in 1-14 Hz range.

| *Model* | *Effects* | *Estimate* ± *SE* | *t* | *p* | *95% CI* | $\sigma^2$ | $\tau_{00}$ | *LL* |
|---|---|---|---|---|---|---|---|---|
| Model s1n | Intercept | 10.47 ± 1.06 | 9.84 | **<.001** | **[8.39 12.38]** | 5.59 | 29.01 | -113.70 |
| Model s1a | Intercept | 1.76 ± 2.88 | 0.61 | .546 | [-3.76 7.76] | 3.47 | 25.23 | -109.29 |
| | MaxPeakFreq | 1.19 ± 0.37 | 3.21 | **.003** | **[0.42 1.92]** | | | |
| Model s1b | Intercept | 10.51 ± 1.06 | 9.93 | **<.001** | **[8.46 12.54]** | 5.93 | 28.11 | -113.64 |
| | Gradient | 1.39 ± 3.63 | 0.38 | .705 | [-6.46 8.62] | | | |
| Model s1c | Intercept | 0.23 ± 2.90 | 0.08 | .938 | [-5.10 6.17] | 4.26 | 19.07 | -106.80 |
| | MaxPeakFreq : Gradient | 1.76 ± 1.08 | 1.63 | .114 | [-0.34 4.07] | | | |

| *Model* | *Effects* | *Estimate* ± *SE* | *z* | *p* | *95% CI* | $\sigma^2$ | $\tau_{00}$ | *LL* |
|---|---|---|---|---|---|---|---|---|
| Model s2n | UWS \| MCS | -2.18 ± 0.91 | -2.39 | **0.017** | **[-3.97 -0.39]** | 1.0 | 9.54 | -73.30 |
| | MCS \| EMCS | -0.13 ± 0.57 | -0.22 | 0.825 | [-1.24 0.99] | | | |
| | EMCS \| HC | 0.86 ± 0.61 | 1.41 | 0.157 | [-0.33 2.056] | | | |
| Model s2a | UWS \| MCS | 15.85 ± 6.92 | 2.29 | **0.022** | **[2.29 29.43]** | 1.0 | 15.58 | -52.43 |
| | MCS \| EMCS | 19.93 ± 8.24 | 2.42 | **0.016** | **[3.78 36.08]** | | | |
| | EMCS \| HC | 22.17 ± 9.16 | 2.42 | **0.016** | **[4.21 40.13]** | | | |
| | MaxPeakFreq | 2.37 ± 0.99 | 2.38 | **0.017** | **[0.42 4.31]** | | | |
| Model s2b | UWS \| MCS | -1.50 ± 0.75 | -2.00 | **0.046** | **[-2.96 -0.03]** | 1.0 | 3.97 | -71.59 |
| | MCS \| EMCS | 0.05 ± 0.40 | 0.13 | 0.900 | [-0.74 0.88] | | | |
| | EMCS \| HC | 0.78 ± 0.46 | 1.71 | 0.088 | [-0.12 1.68] | | | |
| | Gradient | 3.35 ± 1.65 | 2.03 | **0.018** | **[0.11 6.59]** | | | |
| Model s2c | UWS \| MCS | 16.27 ± 6.87 | 2.37 | **0.011** | **[2.80 29.74]** | 1.0 | 14.44 | -47.33 |
| | MCS \| EMCS | 20.42 ± 7.98 | 2.56 | **0.011** | **[4.77 36.07]** | | | |
| | EMCS \| HC | 22.86 ± 8.99 | 2.54 | **0.013** | **[5.25 40.48]** | | | |
| | MaxPeakFreq : Gradient | 2.87 ± 1.64 | 1.74 | 0.081 | [-0.35 6.10] | | | |

*LL* Log-Likelihood, $\sigma^2$ variance of level-1 residual errors, $\tau_{00}$ variance of level-2 residual errors.

**Table B.3.** Results describing *CRSscore* and *CRSdiagnosis* models in a group of patients with multiple maximal peaks in the 1-14 Hz range.

| *Model* | *Effects* | *Estimate* ± *SE* | *t* | *p* | *95% CI* | $\sigma^2$ | $\tau_{00}$ | *LL* |
|---|---|---|---|---|---|---|---|---|
| Model m1n | Intercept | 11.21 ± 1.05 | 10.62 | <**0.001** | **[9.04 13.55]** | 5.81 | 32.21 | -148.05 |
| Model m1a | Intercept | 4.58 ± 2.65 | 1.73 | 0.090 | [-0.89 9.42] | 6.59 | 23.62 | -145.02 |
| | MaxPeakFreq | 0.89 ± 0.33 | 2.68 | **0.010** | **[0.23 1.60]** | | | |
| Model m1b | Intercept | 11.24 ± 1.06 | 10.55 | <**0.001** | **[9.26 13.40]** | 5.85 | 32.01 | -148.03 |
| | Gradient | -0.89 ± 5.16 | -0.17 | 0.864 | [-10.54 9.33] | | | |
| Model m1c | Intercept | -0.51 ± 3.51 | -0.14 | 0.886 | [-7.53 7.45] | 5.87 | 22.09 | -142.85 |
| | MaxPeakFreq : Gradient | -4.16 ± 2.21 | -1.88 | 0.067 | [-8.49 0.52] | | | |

| *Model* | *Effects* | *Estimate* ± *SE* | *z* | *p* | *95% CI* | $\sigma^2$ | $\tau_{00}$ | *LL* | |
|---|---|---|---|---|---|---|---|---|---|
| Model m2n | UWS \| MCS | -2.98 ± 1.03 | -2.89 | **0.004** | **[-5.00 -0.96]** | 1.0 | 13.71 | -80.51 | |
| | MCS \| EMCS | -0.14 ± 0.67 | -0.21 | 0.830 | [-1.46 1.17] | | | | 11 |
| | EMCS \| HC | 1.90 ± 0.81 | 2.34 | **0.019** | **[0.31 3.50]** | | | | |
| Model m2a | UWS \| MCS | 3.65 ± 1.34 | 2.72 | **0.007** | **[1.02 6.29]** | 1.0 | 3.34 | -70.58 | |
| | MCS \| EMCS | 5.86 ± 1.70 | 3.46 | **0.001** | **[2.54 9.19]** | | | | |
| | EMCS \| HC | 7.49 ± 2.06 | 3.63 | <**0.001** | **[3.45 11.53]** | | | | |
| | MaxPeakFreq | 0.71 ± 0.20 | 3.51 | <**0.001** | **[0.31 1.11]** | | | | |
| Model m2b | UWS \| MCS | -3.03 ± 1.05 | -2.89 | **0.004** | **[-5.08 -0.98]** | 1.0 | 13.74 | -80.41 | |
| | MCS \| EMCS | -0.18 ± 0.68 | -0.26 | 0.789 | [-1.51 1.15] | | | | |
| | EMCS \| HC | 1.88 ± 0.81 | 2.31 | **0.021** | **[0.28 3.48]** | | | | |
| | Gradient | -1.22 ± 2.87 | -0.43 | 0.670 | [-6.86 4.40] | | | | |
| Model m2c | UWS \| MCS | 5.16 ± 1.89 | 2.73 | **0.006** | **[1.46 8.86]** | 1.0 | 4.21 | -69.21 | |
| | MCS \| EMCS | 7.58 ± 2.32 | 3.27 | **0.001** | **[3.04 12.12]** | | | | |
| | EMCS \| HC | 9.36 ± 2.71 | 3.45 | **0.001** | **[4.04 14.68]** | | | | |
| | MaxPeakFreq : Gradient | -1.56 ± 1.21 | -1.28 | 0.200 | [-3.94 0.82] | | | | |

*LL* Log-Likelihood, $\sigma^2$ variance of level-1 residual errors, $\tau_{00}$ variance of level-2 residual errors.

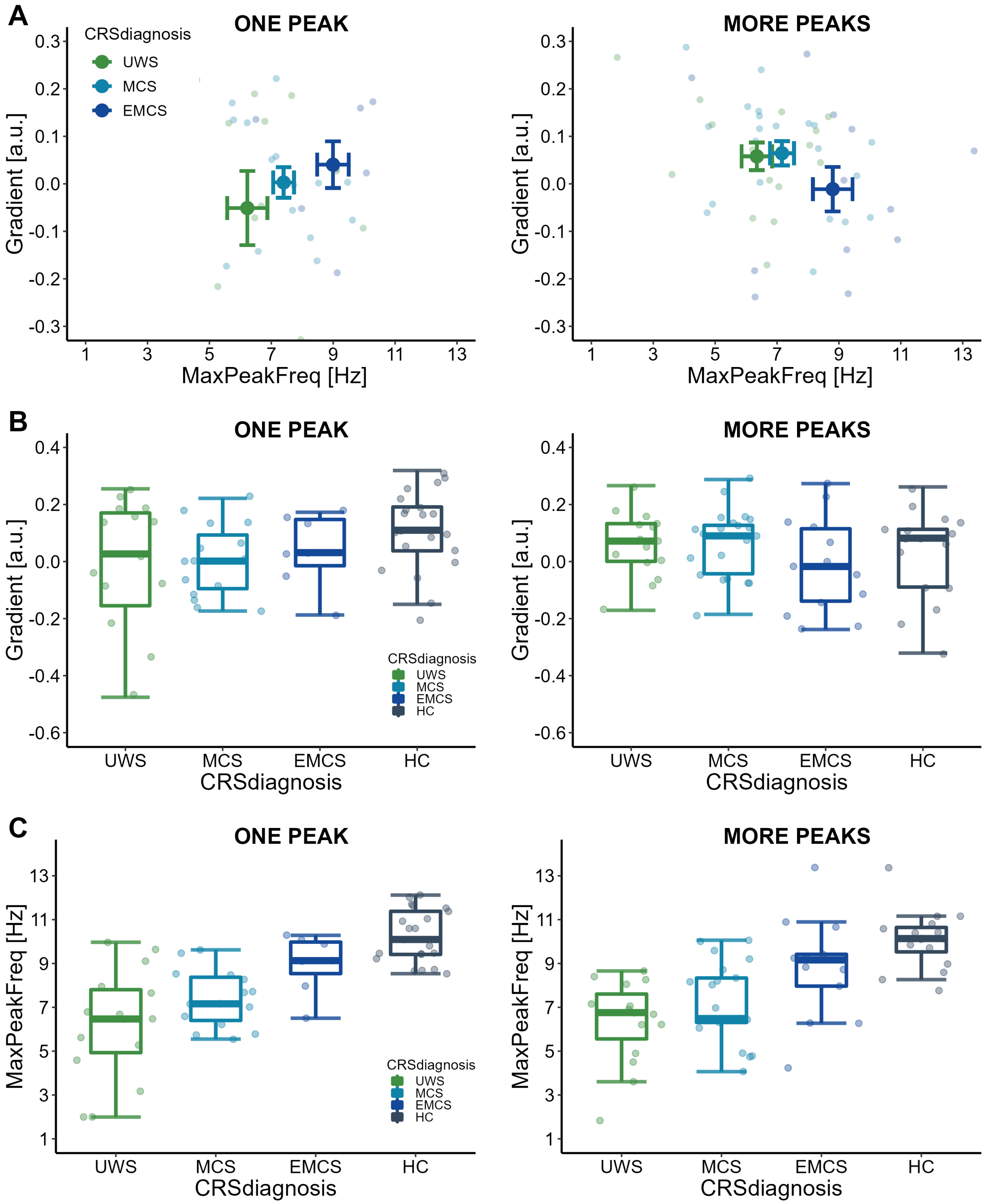


**Figure B.1.** Changes in relationship between MaxPeakFreq, Gradient, CRSscore and CRSdiagnosis in a group of patients with single (*left panels*) or multiple (*right panels*) maximal peaks in 1-14 Hz range. **A** Relationship between *MaxPeakFreq* and *Gradient*. *Dark coloured dots* represent mean value for UWS, MCS and EMCS groups. *Whiskers* represent 95% SE intervals. **B** Values of *Gradient* for different *CRSdiagnosis*. Coloured horizontal bars represent the mean, whiskers represent ± 1.5 * IQR. **C** Values of *MaxPeakFreq* for different *CRSdiagnosis*. Coloured horizontal bars represent the mean, whiskers represent ± 1.5 * IQR.

### Appendix C

#### Controlling for the influence of etiology on the markers of PDOC patients state.

To account for the possibility of relation between measured EEG characteristics and patient's etiology (Forgacs et al 2020; Schiff 2010), we selected a group of 51 patients (80 measurements), who were classified to one of the following etiologies: anoxia, stroke and trauma (demographic statistics in Table **C.1**, Supplementary information). None of the analyses revealed significant differences between *Etiology* groups for *CRSscore* and *CRSdiagnosis* variables (see *Model 1aE* and *Model 2aE* in Table **C.2**, Supplementary information). Furthermore, none of the etiologies improved the relation between both variables and the *MaxpeakFreq* (see *Model 1cE* and *Model 2cE* in Table **C.2**, Supplementary information) or *Gradient* (see *Model 1cE* and *Model 2cE* in Table **C.3**, Supplementary information). We therefore ruled out the possible relation of patients etiology and the improvement of a neurocognitive state in our research (Forgacs et al 2020).

**Table C.1.** Demographics of patient groups within chosen *Etiology*.

| Etiology | | | | | |
|---|---|---|---|---|---|
| *Groups* | *N measurements* | *N* | *Females* | *Age* | |
| | | | | *M* | *SD* |
| anoxia | 14 | 9 | 0 | 42.86 | 14.06 |
| stroke | 13 | 9 | 8 | 43.75 | 13.45 |
| trauma | 53 | 33 | 11 | 33.82 | 13.10 |

**Table C.2.** *CRSscore* and *CRSdiagnosis* models with *MaxPeakFreq* and *Etiology*.

| *Model* | *Effects* | *Estimate* ± *SE* | *t* | *p* | *95% CI* | $\sigma^2$ | $\tau_{00}$ | *LL* |
|---|---|---|---|---|---|---|---|---|
| Model 1nE | Intercept | 10.72 ± 0.83 | 12.94 | **<0.001** | **[9.09 12.29]** | 4.85 | 31.34 | -235.76 |
| Model 1aE | trauma : stroke | -0.17 ± 2.21 | -0.08 | 0.997 | [-5.50 5.16] | 4.81 | 28.82 | -233.71 |
| | trauma : anoxia | 4.28 ± 2.21 | 1.94 | 0.138 | [-1.04 9.60] | | | |
| | stroke : anoxia | 4.45 ± 2.77 | 1.60 | 0.252 | [-2.23 11.12] | | | |
| Model 1bE | Intercept | 5.53 ± 1.82 | 3.03 | **<0.003** | **[2.01 9.45]** | 4.74 | 26.10 | -231.22 |
| | MaxPeakFreq | 0.71 ± 0.23 | 3.14 | **<0.003** | **[0.21 1.14]** | | | |
| Model 1cE | MaxPeakFreq : trauma | 0.76 ± 0.43 | 1.78 | 0.078 | [-0.09 1.62] | 4.69 | 24.58 | -229.71 |
| | MaxPeakFreq : stroke | 0.62 ± 0.47 | 1.32 | 0.194 | [-0.33 1.57] | | | |
| | MaxPeakFreq : anoxia | 0.60 ± 0.37 | 1.63 | 0.109 | [-0.14 1.34] | | | |

| *Model* | *Effects* | *Estimate* ± *SE* | *z* | *p* | *95% CI* | $\sigma^2$ | $\tau_{00}$ | *LL* | |
|---|---|---|---|---|---|---|---|---|---|
| Model 2nE | UWS \| MCS | -0.80 ± 0.54 | -1.47 | 0.143 | [-1.86 0.27] | 1.0 | 8.13 | -74.12 | 25 |
| | MCS \| EMCS | 2.41 ± 0.88 | 2.73 | **0.006** | **[0.68 4.15]** | | | | |
| Model 2aE | UWS \| MCS | 3.50 ± 1.44 | 2.43 | **0.015** | **[0.68 6.32]** | 1.0 | 5.24 | -67.59 | |
| | MCS \| EMCS | 6.54 ± 1.91 | 3.43 | **0.008** | **[2.80 10.28]** | | | | |
| | MaxPeakFreq | 0.58 ± 0.20 | 2.96 | **0.003** | **[0.20 0.97]** | | | | |
| Model 2bE | UWS \| MCS | -1.08 ± 0.59 | -1.82 | 0.069 | [-2.24 0.08] | 1.0 | 6.19 | -70.98 | |
| | MCS \| EMCS | 1.97 ± 0.74 | 2.65 | **0.008** | **[0.51 3.42]** | | | | |
| | trauma : stroke | -0.79 ± 1.10 | -0.72 | 0.753 | [-3.36 1.79] | | | | |
| | trauma : anoxia | 2.71 ± 1.36 | 1.99 | 0.114 | [-0.48 5.91] | | | | |
| | stroke : stroke | 3.50 ± 1.67 | 2.10 | 0.090 | [-0.41 7.41] | | | | |
| Model 2cE | UWS \| MCS | 1.47 ± 1.74 | 0.84 | 0.399 | [-1.95 4.89] | 1.0 | 5.51 | -61.79 | |
| | MCS \| EMCS | 4.86 ± 1.95 | 2.50 | **0.013** | **[1.04 8.68** | | | | |
| | MaxPeakFreq : trauma | 0.37 ± 0.23 | 1.60 | 0.109 | [-0.08 0.82] | | | | |
| | MaxPeakFreq : stroke | 2.42 ± 1.12 | 2.16 | **0.030** | **[0.23 4.62]** | | | | |
| | MaxPeakFreq : anoxia | 0.73 ± 0.43 | 1.70 | 0.089 | [-0.11 1.58] | | | | |

*LL* Log-Likelihood, $\sigma^2$ variance of level-1 residual errors, $\tau_{00}$ variance of level-2 residual errors.

**Table C.3.** *CRSscore* and *CRSdiagnosis* models with *Gradient* and *Etiology*.

| *Model* | *Effects* | *Estimate* ± *SE* | *t* | *p* | *95% CI* | $\sigma^2$ | $\tau_{00}$ | *LL* |
|---|---|---|---|---|---|---|---|---|
| Model 1nE | Intercept | 10.72 ± 0.83 | 12.94 | <**0.001** | **[9.07 12.28]** | 4.85 | 31.34 | -235.76 |
| Model 1aE | trauma : stroke | -0.17 ± 2.21 | -0.08 | 0.997 | [-5.50 5.16] | 4.81 | 28.82 | -233.71 |
| | trauma : anoxia | 4.28 ± 2.21 | 1.94 | 0.138 | [-1.04 9.60] | | | |
| | stroke : anoxia | 4.45 ± 2.77 | 1.60 | 0.252 | [-2.23 11.12] | | | |
| Model 1bE | Intercept | 10.72 ± 0.83 | 12.92 | <**0.001** | **[9.09 12.37]** | 4.84 | 31.40 | -235.76 |
| | Gradient | -0.31 ± 2.61 | -0.12 | 0.905 | [-5.79 4.94] | | | |
| Model 1cE | Gradient : trauma | -2.52 ± 4.37 | -0.58 | 0.567 | [-11.29 6.25] | 4.77 | 28.23 | -233.09 |
| | Gradient : stroke | -4.64 ± 6.34 | -0.73 | 0.469 | [-17.45 8.18] | | | |
| | Gradient : anoxia | 2.13 ± 4.24 | 0.50 | 0.618 | [-6.33 10.58] | | | |

| *Model* | *Effects* | *Estimate* ± *SE* | *z* | *p* | *95% CI* | $\sigma^2$ | $\tau_{00}$ | *LL* | |
|---|---|---|---|---|---|---|---|---|---|
| Model 2nE | UWS \| MCS | -0.76 ± 0.49 | -1.53 | 0.125 | [-1.73 0.21] | 1.00 | 6.86 | -74.34 | 26 |
| | MCS \| EMCS | 2.25 ± 0.75 | 2.99 | **0.003** | **[0.77 3.72]** | | | | |
| Model 2aE | UWS \| MCS | -0.76 ± 0.50 | -1.53 | 0.126 | [-1.73 0.21] | 1.00 | 6.95 | -74.33 | |
| | MCS \| EMCS | 2.26 ± 0.76 | 2.97 | **0.003** | **[0.77 3.75]** | | | | |
| | Gradient | -0.22 ± 1.79 | -0.12 | 0.901 | [-3.72 3.28] | | | | |
| Model 2bE | UWS \| MCS | -1.08 ± 0.59 | -1.82 | 0.069 | [-2.24 0.08] | 1.00 | 6.19 | -70.98 | |
| | MCS \| EMCS | 1.97 ± 0.74 | 2.65 | <**0.008** | **0.51 3.42]** | | | | |
| | trauma : stroke | -0.79 ± 1.10 | -0.72 | 0.753 | [-3.36 1.79] | | | | |
| | trauma : anoxia | 2.71 ± 1.36 | 1.99 | 0.114 | [-0.48 5.91] | | | | |
| | stroke : stroke | 3.50 ± 1.67 | 2.10 | 0.090 | [-0.41 7.41] | | | | |
| Model 2cE | UWS \| MCS | -1.17 ± 0.63 | -1.86 | 0.062 | [-2.40 0.06] | 1.00 | 6.95 | -68.66 | |
| | MCS \| EMCS | 2.08 ± 0.78 | 2.66 | <**0.008** | **[0.54 3.61]** | | | | |
| | Gradient : trauma | -1.35 ± 2.86 | -0.47 | 0.638 | [-6.95 4.25] | | | | |
| | Gradient : stroke | -8.73 ± 6.56 | -1.33 | 0.183 | [-21.58 4.12] | | | | |
| | Gradient : anoxia | 4.97 ± 4.51 | 1.10 | 0.270 | [-3.87 13.82] | | | | |

*LL* Log-Likelihood, $\sigma^2$ variance of level-1 residual errors, $\tau_{00}$ variance of level-2 residual errors.

#### Appendix D

##### Controlling for the influence of drug intake on the markers of PDOC patients state.

To account for the possibility of relation between measured EEG characteristics and received medication (Depakine, Zenkov 2002; Amantix, Terzano et al 1983; Baclofen, Badr et al 1983; Ciurleo et al 2013) we selected a group of 36 patients (46 measurements), for whose this information was available: Amantix, Baclofen, Depakine, and finally no medication of interest. (demographic statistics in Table D.1, Supplementary information).

None of the analyses revealed significant differences between *Medications* groups for *CRSscore* and *CRSdiagnosis* variables (see *Model 1aM* and *Model 2aM* in Table D.2, Supplementary information). Furthermore, none of the etiologies improved the relation between both variables and the *MaxpeakFreq* (see *Model 1cM* and *Model 2cM* in Table D.2, Supplementary information) or *Gradient* (see *Model 1cM* and *Model 2cM* in Table D.3, Supplementary information). Therefore, in our opinion, drug intake did not serve as an interfering factor.

**Table D.1.** Demographic features of groups of patients within chosen *Medication*.

| Medications | | | | | |
|---|---|---|---|---|---|
| *Groups* | *N measurements* | *N* | *Females* | *Age* | |
| | | | | *M* | *SD* |
| no drugs | 10 | 7 | 3 | 40.83 | 14.39 |
| Amantix | 7 | 4 | 2 | 37.71 | 12.45 |
| Baclofen | 20 | 16 | 3 | 38.95 | 12.29 |
| Depakine | 9 | 9 | 3 | 33.78 | 10.54 |

**Table D.2.** *CRSscore* and *CRSdiagnosis* models with *MaxPeakFreq* and *Medication*.

| *Model* | *Effects* | *Estimate* ± *SE* | *t* | *p* | *95% CI* | $\sigma^2$ | $\tau_{00}$ | *LL* |
|---|---|---|---|---|---|---|---|---|
| Model 1nM | Intercept | 9.92 ± 1.18 | 8.4 | **<0.001** | **[7.43 12.57]** | 4.83 | 24.58 | -101.58 |
| Model 1aM | Intercept | -2.37 ± 2.71 | -0.87 | 0.388 | [-7.40 2.95] | 2.11 | 22.15 | -93.29 |
| | MaxPeakFreq | 1.72 ± 0.35 | 4.93 | **<0.001** | **[1.00 2.36]** | | | |
| Model 1bM | Amantix : Baclofen | 0.23 ± 1.39 | 0.17 | 0.985 | [-3.28 3.74] | 4.78 | 24.64 | -101.51 |
| | Amantix : Depakine | -0.16 ± 1.72 | -0.10 | 0.995 | [-4.49 4.16] | | | |
| | Baclofen : Depakine | -0.40 ± 1.24 | -0.32 | 0.995 | [-3.51 2.71] | | | |
| Model 1cM | MaxPeakFreq : Amantix | 2.32 ± 0.69 | 3.36 | **0.003** | **[0.89 3.75]** | 1.86 | 22.14 | -92.22 |
| | MaxPeakFreq : Baclofen | 1.40 ± 0.49 | 2.87 | **0.006** | **[0.41 2.39]** | | | |
| | MaxPeakFreq : Depakine | 1.63 ± 0.60 | 2.73 | **0.009** | **[0.42 2.83]** | | | |
| *Model* | *Effects* | *Estimate* ± *SE* | *z* | *p* | *95% CI* | $\sigma^2$ | $\tau_{00}$ | *LL* |
| Model 2nM | UWS \| MCS | -0.85 ± 0.93 | -0.92 | 0.357 | [-2.67 0.96] | 1.0 | 10.44 | -30.36 |
| | MCS \| EMCS | 3.52 ± 1.69 | 2.08 | **0.037** | **[0.21 6.83]** | | | |
| Model 2aM | model did not converge | | | | | | | |
| Model 2bM | UWS \| MCS | -0.92 ± 1.15 | -0.80 | 0.423 | [-3.17 1.33] | 1.0 | 10.39 | -30.35 |
| | MCS \| EMCS | 3.45 ± 1.99 | 1.73 | 0.084 | [-0.46 7.35] | | | |
| | Amantix : Baclofen | 0.09 ± 0.81 | 0.11 | 0.993 | [-1.80 1.98] | | | |
| | Amantix : Depakine | 0.04 ± 1.18 | 0.04 | 0.999 | [-2.73 2.81] | | | |
| | Baclofen : Depakine | -0.05 ± 1.00 | -0.05 | 0.999 | [-2.39 2.29] | | | |
| Model 2cM | model did not converge | | | | | | | |

*LL* Log-Likelihood, $\sigma^2$ variance of level-1 residual errors, $\tau_{00}$ variance of level-2 residual errors.

**Table D.3.** *CRSscore* and *CRSdiagnosis* models with *Gradient* and *Medication*.

| *Model* | *Effects* | *Estimate* ± *SE* | *t* | *p* | *95% CI* | $\sigma^2$ | $\tau_{00}$ | *LL* |
|---|---|---|---|---|---|---|---|---|
| Model 1nM | Intercept | 9.92 ± 1.18 | 8.40 | **<.001** | **[7.47 12.03]** | 4.83 | 24.58 | -101.58 |
| Model 1aM | Intercept | 10.00 ± 1.16 | 8.59 | **<.001** | **[7.71 12.42]** | 4.78 | 24.64 | -101.32 |
| | Gradient | 3.15 ± 4.32 | 0.73 | .472 | [-5.79 12.67] | | | |
| Model 1bM | Amantix : Baclofen | 0.23 ± 1.39 | 0.17 | .985 | [-3.28 3.74] | 4.89 | 23.52 | -101.51 |
| | Amantix : Depakine | -0.16 ± 1.72 | -0.10 | .995 | [-4.49 4.16] | | | |
| | Baclofen : Depakine | -0.40 ± 1.24 | -0.32 | .945 | [-3.51 2.71] | | | |
| Model 1cM | Gradient : Amantix | 3.56 ± 10.61 | 0.34 | .740 | [-18.38 25.5] | 4.88 | 23.35 | -101.24 |
| | Gradient : Baclofen | 3.21 ± 5.17 | 0.62 | .538 | [-7.27 13.7] | | | |
| | Gradient : Depakine | 1.76 ± 7.50 | 0.23 | .815 | [-13.37 16.9] | | | |

| *Model* | *Effects* | *Estimate* ± *SE* | *z* | *p* | *95% CI* | $\sigma^2$ | $\tau_{00}$ | *LL* |
|---|---|---|---|---|---|---|---|---|
| Model 2nM | UWS \| MCS | -0.85 ± 0.93 | -0.92 | 0.357 | [-2.67 0.96] | 1.00 | 10.44 | -30.36 |
| | MCS \| EMCS | 3.52 ± 1.69 | 2.08 | **0.037** | **[0.21 6.83]** | | | |
| Model 2aM | UWS \| MCS | -0.86 ± 0.87 | -0.99 | 0.323 | [-2.56 0.84] | 1.00 | 7.85 | -29.99 |
| | MCS \| EMCS | 3.15 ± 1.60 | 1.97 | **0.049** | **[0.01 6.30]** | | | |
| | Gradient | 2.62 ± 2.92 | 0.90 | 0.370 | [-3.11 8.34] | | | |
| Model 2bM | UWS \| MCS | -0.92 ± 1.15 | -0.80 | 0.423 | [-3.17 1.33] | 1.00 | 10.39 | -30.35 |
| | MCS \| EMCS | 3.45 ± 1.99 | 1.73 | 0.084 | [-0.46 7.35] | | | |
| | Amantix : Baclofen | 0.09 ± 0.81 | 0.11 | 0.993 | [-1.80 1.98] | | | |
| | Amantix : Depakine | 0.04 ± 1.18 | 0.04 | 0.999 | [-2.73 2.81] | | | |
| | Baclofen : Depakine | -0.05 ± 1.00 | -0.05 | 0.999 | [-2.39 2.29] | | | |
| Model 2cM | UWS \| MCS | -1.03 ± 1.02 | -1.01 | 0.311 | [-3.04 0.97] | 1.00 | 6.27 | -29.66 |
| | MCS \| EMCS | 2.71 ± 1.89 | 1.44 | 0.150 | [-0.98 6.41] | | | |
| | Gradient : Amantix | 0.20 ± 5.65 | 0.04 | 0.971 | [-10.88 11.3] | | | |
| | Gradient : Baclofen | 3.78 ± 3.38 | 1.12 | 0.262 | [-2.83 10.4] | | | |
| | Gradient : Depakine | 0.78 ± 5.12 | 0.15 | 0.879 | [-9.27 10.8] | | | |

*LL* Log-Likelihood, $\sigma^2$ variance of level-1 residual errors, $\tau_{00}$ variance of level-2 residual errors.
